# Cell type profiling in salamanders identifies innovations in vertebrate forebrain evolution

**DOI:** 10.1101/2022.03.28.485354

**Authors:** Jamie Woych, Alonso Ortega Gurrola, Astrid Deryckere, Eliza C. B. Jaeger, Elias Gumnit, Gianluca Merello, Jiacheng Gu, Alberto Joven Araus, Nicholas D. Leigh, Maximina Yun, András Simon, Maria Antonietta Tosches

**Affiliations:** Department of Biological Sciences, Columbia University; New York City, 10027 New York, USA; Department of Neuroscience, Columbia University; New York City, 10027 New York, USA; Department of Cell and Molecular Biology, Karolinska Institute; Stockholm, Sweden; Molecular Medicine & Gene Therapy, Wallenberg Centre for Molecular Medicine, Lund Stem Cell Center; Lund, Sweden; Technische Universität Dresden, CRTD/Center for Regenerative Therapies Dresden; Dresden, Germany; Max Planck Institute for Molecular Cell Biology and Genetics; Dresden, Germany

## Abstract

The evolution of advanced cognition in vertebrates is associated with two independent innovations in the forebrain: the six-layered neocortex in mammals and the dorsal ventricular ridge (DVR) in sauropsids (reptiles and birds). How these novelties arose in vertebrate ancestors remains unclear. To reconstruct forebrain evolution in tetrapods, we built a cell type atlas of the telencephalon of the salamander *Pleurodeles waltl*. Our molecular, developmental, and connectivity data indicate that parts of the sauropsid DVR trace back to tetrapod ancestors. In contrast, the salamander dorsal pallium is devoid of cellular and molecular characteristics of the mammalian neocortex, yet shares similarities with entorhinal cortex and subiculum. Our findings chart the series of innovations that resulted in the emergence of the sauropsid DVR, and the mammalian six-layered neocortex.

## Introduction

The transition from water to land was a pivotal moment in vertebrate history that exposed the first tetrapods to new environmental and cognitive challenges, which may have accelerated adaptive innovations in the nervous system (1). After the divergence of mammals and sauropsids (reptiles and birds) about ∼320 million years ago, innovations in the pallium (i.e. dorsal telencephalon) paved the way for advanced cognition. In mammals, the neocortex, with its characteristic six-layers, evolved from a simpler ancestral cortex located in the dorsal pallium (2). In sauropsids, an expansion of the ventral pallium produced a large set of nuclei called the dorsal ventricular ridge (DVR) (Fig. 1A). Although neocortex and DVR develop from different parts of the pallium, they bear extensive similarities in terms of gene expression, connectivity, and function (3–5). A model centered on brain connectivity proposes the homology of neocortex and DVR, implying that differences in neocortex and DVR development and topological positions arose secondarily (6). Developmental studies (2) and recent adult transcriptomics data (7, 8) challenge this view, suggesting that DVR and neocortex have separate evolutionary origins in amniote ancestors, and therefore similar functions were acquired independently. However, the innovations that led to the DVR and neocortex remain poorly understood at the molecular and cellular levels.

**Figure 1:**
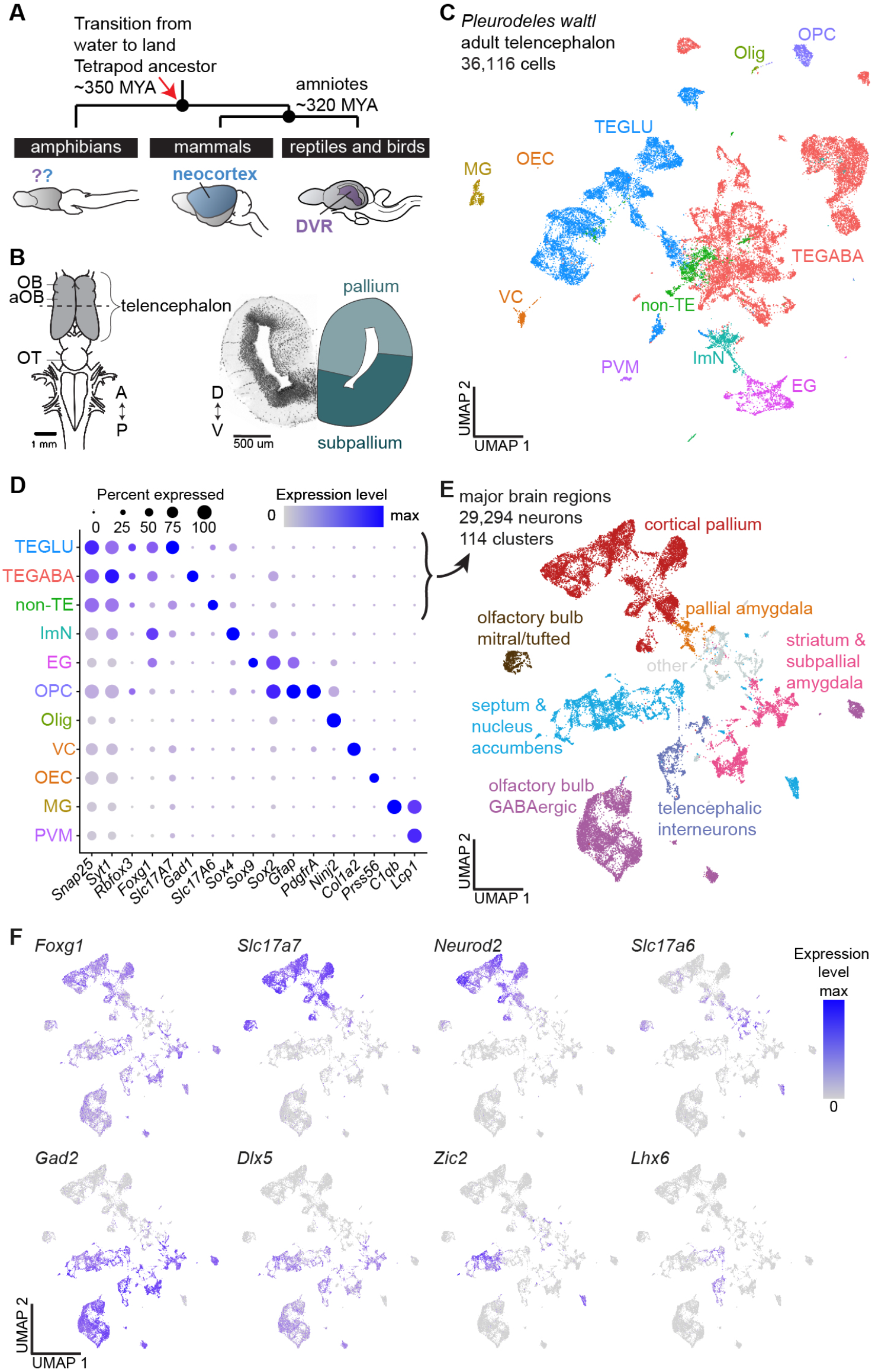
Neuronal diversity in the *Pleurodeles waltl* telencephalon. **(A)** Schematic highlighting the phylogenetic position of amphibians, the mammalian neocortex, and the reptilian DVR. **(B)** Left: schematic of the *Pleurodeles waltl* brain (dorsal view). Dotted line indicates section plane for coronal slice on the right. **(C)** UMAP (Uniform Manifold Approximation and Projection) plot of 36,116 salamander single-cell transcriptomes, colors indicate cell classes. **(D)** Dot plot showing the expression of marker genes used to annotate the telencephalic dataset in **(C). (E)** UMAP plot of 29,294 single-cell transcriptomes of salamander neurons, colors indicate major brain regions. **(F)** UMAPs showing expression of key markers of glutamatergic and GABAergic neurons in the neuronal dataset. Abbreviations: A, anterior; aOB, accessory olfactory bulb; D, dorsal; EG, ependymal glia; GLU, glutamatergic; ImN, immature neurons; MG, microglia; MYA, million years ago; OB, olfactory bulb; OEC, olfactory ensheathing cells; Olig, oligodendrocytes; OPC, oligodendrocyte precursor cells; OT, optic tectum; P, posterior; PVM, perivascular macrophages; TE, telencephalic; V, ventral; VC, vascular cells

We reasoned that if neocortex and DVR have separate origins, they may trace back to pallial regions that existed in a pre-amniote ancestor. Amphibians, who diverged from the rest of tetrapods ∼350 million years ago, have a simple telencephalic architecture, devoid of obvious layering or large brain nuclei (Fig. 1B). Both a dorsal and a ventral pallium exist in amphibians (9), but it is unclear whether they are related in any way to neocortex and DVR. Here, we analyzed the telencephalon of *Pleurodeles waltl*, a salamander species with a true adult (post-metamorphic) stage, to ask the following: (i) Are there neuron types in the amphibian pallium with transcriptomic similarity to neocortical or DVR neurons? (ii) How do these neurons develop? (iii) Do these neurons display patterns of connectivity similar to neocortex or DVR?

## Results

### A cell type atlas of the salamander telencephalon

To build a cell type atlas of the salamander telencephalon, we profiled entire brains and microdissected telencephali of adult *Pleurodeles* (brain atlas in Fig. S1, Movie S1). After single-cell RNA sequencing (scRNAseq, 10x Genomics), reads were mapped on a new long-read de novo reference transcriptome (see Methods). Following quality filtering, we obtained 36,116 single-cell transcriptomes, performed Louvain clustering, and identified 11 major populations of neuronal and non-neuronal cells (Fig. 1C; Fig. S2).

We annotated clusters of differentiated neurons, immature neurons, ependymoglial cells, microglia, oligodendrocytes, oligodendrocyte precursors, and other non-neuronal cells based on well-established marker genes (Fig. 1D, Fig. S2C). Differentiated neurons (29,294 cells), identified by the expression of pan-neuronal markers such as *Snap25, Syt1*, and *Rbfox3* (i.e., *NeuN*), were subclustered to classify neuron types. This revealed 47 clusters of glutamatergic neurons and 67 clusters of GABAergic neurons, which we annotated on the basis of marker genes with conserved expression across species, *in situ* hybridization for newly-identified markers, and existing amphibian literature (reviewed in (10, 11)) (Fig. 1E,F; Figs. S3-6).

In the telencephalon, hierarchical clustering revealed four distinct groups of glutamatergic clusters (Fig. S3A). One expressed *Neurod2* and *Slc17a7* (*Vglut1*) at high levels; we named this group cortical pallium for its molecular similarity to the cerebral cortex of mammals and reptiles (12, 13). The remaining groups included olfactory bulb mitral and tufted cells, expressing the transcription factor *Tbx21* (14); pallial amygdala (pA) neurons, expressing lower levels of *Slc17a7* and *Neurod2* and high levels of *Slc17a6* (*Vglut2*); and glutamatergic neurons in the septum, expressing *Slc17a6, Zic2*, and *Isl1* (Fig. 1E,F; Fig. S3A).

Telencephalic GABAergic neurons express markers of the subpallium such as *Dlx5, Gad1* and *Gad2*. We found that the amphibian subpallium includes not only neurons from lateral and medial ganglionic eminences (LGE and MGE) as previously shown (15, 16), but also from the caudal ganglionic eminence (CGE). Specifically, we identified several types of striatal and septal neurons, nucleus accumbens, bed nucleus of the stria terminalis, and diagonal band neurons, as well as olfactory bulb LGE-derived GABAergic interneurons (Fig. S3A; Fig. S4A-C). Telencephalic GABAergic interneurons, scattered throughout the pallium, included MGE- and CGE-derived cells (Fig. S4F). Despite its anatomical simplicity, these data indicate that the amphibian telencephalon harbors a greater degree of cell-type complexity than anticipated.

### Regional and layered distribution of pallial glutamatergic neurons

Previous literature indicates that the amphibian pallium is organized in medial, dorsal, lateral, and ventral pallium (MP, DP, LP, and VP) (9, 17), but precise boundaries and further subdivisions are a matter of dispute (18, 19). To clarify the organization of the amphibian telencephalon, we built a transcriptomics-based map in *Pleurodeles*.

Hierarchical clustering of average cluster expression profiles indicates a clear distinction between cortical pallium and pallial amygdala (pA), and the existence of four groups of cortical pallium clusters (Fig. 2A,B, Fig. S3A,B). As shown by *in situ* hybridization for specific marker genes (Fig. 2C), the four groups largely correspond to MP, DP, LP, and VP. In mid-telencephalic sections, the MP, classically compared to the hippocampus for its position and connectivity, expressed hippocampal transcription factors such as *Fezf2, Lhx9, Zbtb20*, and *Etv1* (7, 20). The DP, anatomically distinct from the MP, expressed low-levels of MP markers but high levels of *Etv1*. The LP, a narrow band of densely-packed neurons, expressed *Lhx2, Satb1, Rorb*, and *Reln*, markers of olfactory-recipient cells in the mammalian piriform cortex (semilunar cells) (21). Most of the ventral pallium expresses the transcription factor Sox6 and is molecularly diverse, in line with its anatomical heterogeneity (9, 22). Subdivisions include an anterior *Znf536*+/*Grm3*+/*Nr2f2*-post-olfactory eminence (POE) (23, 24), a *Nos1*-anterior VP (VPa), and a *Nos1*+ posterior VP (VPp), see also (16). Along the anterior-posterior axis, *Slc17a6* is expressed only in the most ventral portion of the VP (Fig. S5,6; Movies S2,3).

**Figure 2:**
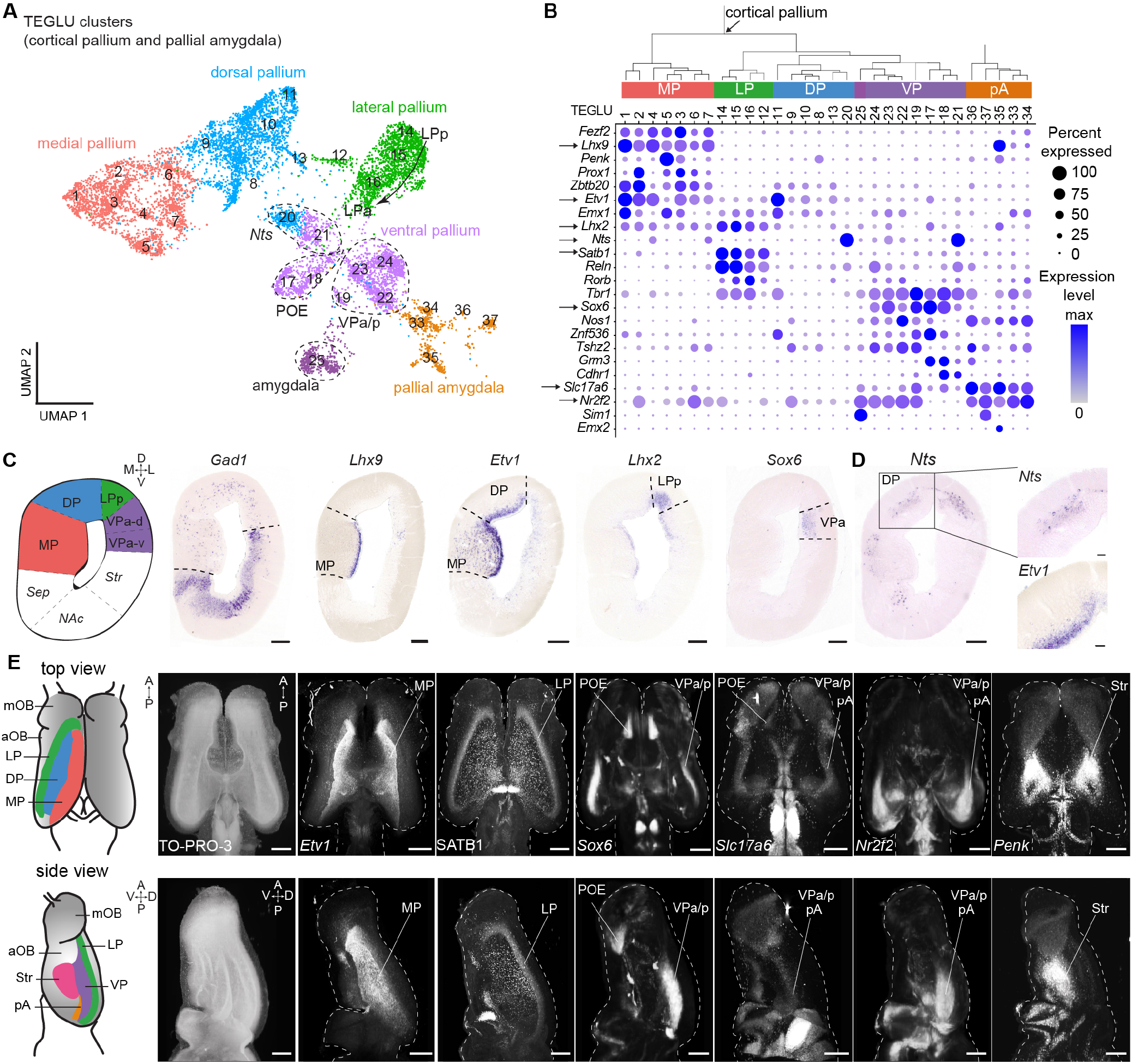
Spatial mapping of pallial neurons in *Pleurodeles*. **(A)** UMAP plot of clusters from cortical pallium and pallial amygdala, annotated by the inferred pallial region. **(B)** Top: Hierarchical clustering of average cluster expression; splits are based on molecular similarity and do not indicate developmental or evolutionary relationships. Bottom: dot plot showing the expression of key marker genes defining distinct pallial regions. Arrows: genes shown in C and E. **(C)** Left to right: schematic of a coronal section at mid-telencephalic level, expression of Gad1, indicating the pallial-subpallial boundary, and of transcription factors labeling distinct pallial regions along the mediolateral axis; Scale bars: 200 um. **(D)** Expression of Nts and Etv1 in layers, boxed areas indicate magnifications on the right. Scale bars in right panels: 50 um. **(E)** Left: schematics of dorsal and lateral surfaces of the salamander telencephalon. Right: dorsal and lateral views of whole-mount immunohistochemistry or HCR stainings for telencephalic markers. Panels show maximum intensity projections of brains after clearing and volumetric light-sheet imaging. Scale bars: 500 um. See methods for specifics on SATB1 antibody.

A closer look at *Nts*, a marker expressed at high levels in two clusters (Fig. 2B), revealed differential expression along the radial axis. Visualization of *Nts* expression in tandem with genes expressed in cells closer to the ventricle (e.g. *Etv1* in MP/DP) clearly demonstrates that *Nts* demarcates a discrete, superficial layer of the pallium (Fig. 2D; Fig. S5H). These results are the first suggesting that the amphibian pallium contains at least two separate layers of distinct neuron types.

To resolve the 3D organization of the pallium, we exploited the small size of the salamander brain to combine whole-mount hybridization chain reaction (HCR) *in situ* hybridization, brain clearing (iDISCO), and light-sheet imaging, creating a 3D molecular map (Fig. 2E; Movies S2,3). This revealed that the cortical pallium is organized in adjacent longitudinal stripes, running the length of the telencephalic vesicle (Fig. 2E). For example, the LP extends from the most rostral tip of the pallium, where it contacts the main olfactory bulb (mOB) and the POE, to the caudal tip of the telencephalon. Instead, the pA is localized caudally and is demarcated by *Slc17a6* and *Nr2f2* expression, and absence of *Sox6* (Fig. 2E; Fig. S6; Movies S2,3). Together, these data represent a transcriptomics-based map of the amphibian pallium, and support the existence of distinct regions along the mediolateral axis and distinct layers along the radial axis.

### Distinct developmental trajectories of dorsomedial and ventrolateral pallium

How regions of the amphibian pallium compare to distinct regions of the mammalian and sauropsid pallium, including hippocampus, olfactory cortex, and amygdala, remains debated (10, 25, 26). Current models postulate that pallial regions are homologous when they develop from homologous progenitor domains (17, 27, 28). To trace the developmental history of *Pleurodeles* pallial neuron types, we collected scRNAseq data from stage 36, 41, and 50 larvae (Fig. 3A, Fig. S7A) (29). After unsupervised clustering, we identified radial glia and telencephalic glutamatergic and GABAergic developing neurons (Fig. 3B,C; Fig. S7B,C). To assign developing neurons to their terminal fate in the adult telencephalon, we mapped adult scRNAseq data on developmental data using the Seurat label transfer algorithm (see Methods) (30). This showed that our larval dataset included differentiating neurons from all major pallial and subpallial subdivisions (Fig. 3D; Fig. S7E-J). We then inferred developmental trajectories with Slingshot (31), identifying two trajectories for the differentiation into olfactory bulb mitral and tufted cells, one trajectory for dorsomedial (DM) pallium, and one for ventrolateral (VL) pallium (pA neurons were excluded because their progenitors were poorly sampled, see Methods) (Fig. 3E; S7K). After reordering cells according to their pseudotime score, we compared gene expression along these trajectories. Transcription factors upregulated along the ventrolateral trajectory included *Pbx3* and *Sox6*, expressed in the developing mouse ventral pallium (32). Transcription factors upregulated along the dorsomedial trajectory included the MP markers *Zbtb20, Lhx9, Fezf2, Etv1*, and *Prox1* (Fig. 2B; Fig. 3F; Fig. S5; Fig. S7J,L), in line with label transfer results. This indicates that neurons in the dorsomedial and ventrolateral pallium are specified by distinct gene regulatory cascades, possibly controlled by medial wnt signaling and ventrolateral wnt antagonists (28). Notably, we did not identify distinct developmental trajectories for MP and DP, or LP and VP, suggesting that during early development these pallial regions are not as sharply defined as proposed previously (32, 33).

**Figure 3:**
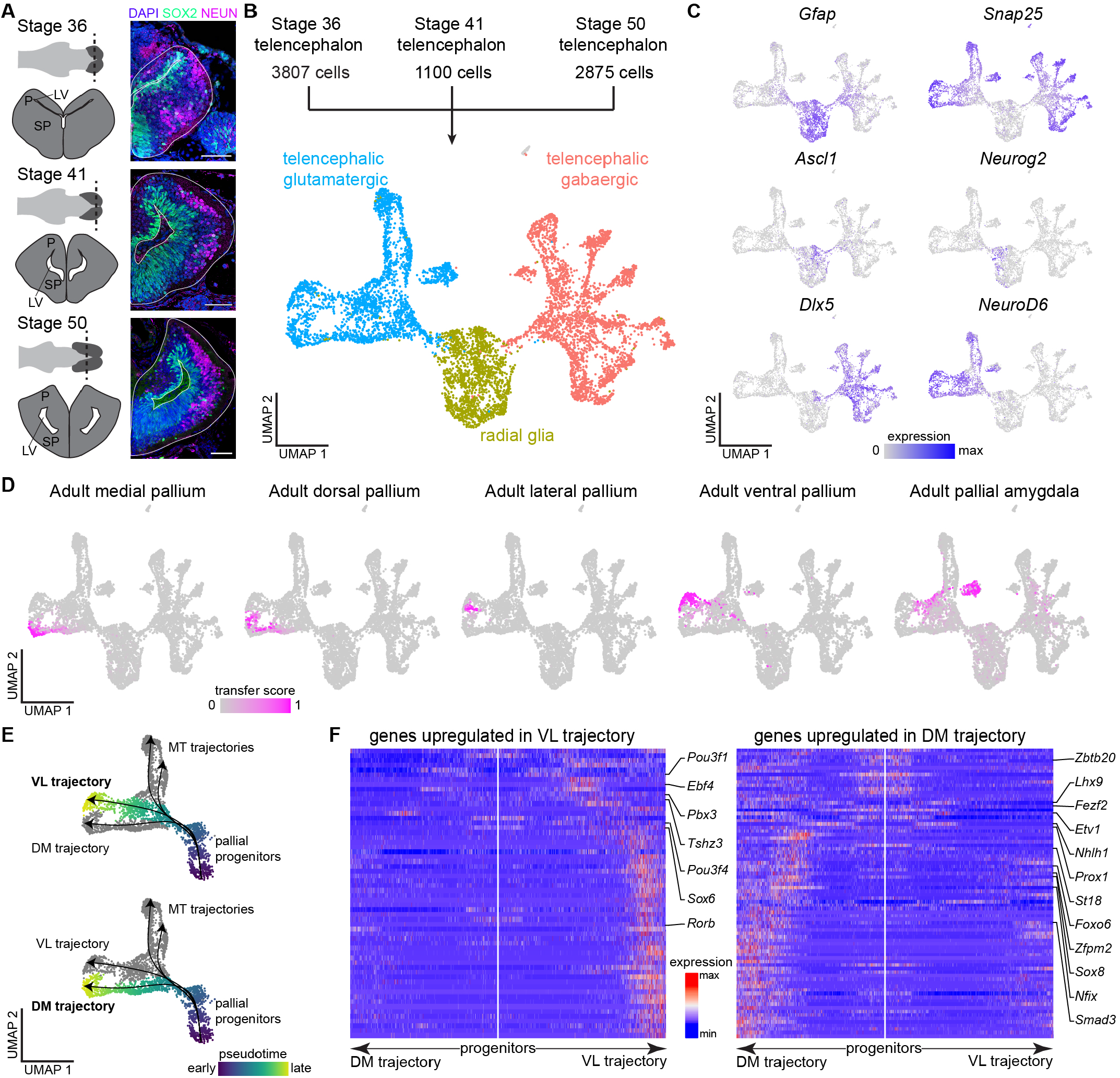
Developmental trajectories in the salamander telencephalon. **(A)** Overview of telencephalic development in *Pleurodeles*. Right: coronal sections through the telencephalon showing radial glia and interneurons stained with SOX2, and differentiated neurons stained with NEUN. Scale bars: 100 um. **(B)** UMAP plot of 7,782 telencephalic cells across the three developmental stages sampled. **(C)** UMAP plot showing single cells colored according to the expression of *Gfap* (radial glia), *Snap25* (differentiated neurons), *Ascl1* and *Neurog2* (committed subpallial and pallial progenitors), and *Dlx5* and *NeuroD6* (postmitotic subpallial and pallial neurons). **(D)** UMAP plots of the developmental dataset, with single cells color-coded according to the label transfer score of adult pallial regions on developmental data. **(E)** Developmental trajectories, ventrolateral or dorsomedial pallial neurons color-coded by their pseudotime score. **(F)** Heatmaps of genes differentially expressed along the differentiation trajectories with transcription factors highlighted on the side. White line in the middle of each panel indicates the position of the pallial progenitors and arrows to the left and right represent the two trajectories. Gene expression levels for each gene are scaled by root mean square ranging from -2 to 6. Abbreviations: DM, dorsomedial; LV, lateral ventricle; MT, mitral and tufted cells of the olfactory bulb; P, pallium; SP, subpallium; VL, ventrolateral.

### Molecular similarity of salamander ventral pallium and reptilian anterior dorsal ventricular ridge

Informed by our results on development, we compared neuron types in the dorsomedial and ventrolateral pallia of salamanders, reptiles, and mammals. To compare scRNAseq data across species, we used a data integration algorithm based on the identification of mutual nearest-neighbors across single-cell datasets (Seurat integration, see Methods) (30). To avoid potential overfitting, we limited data integration to single cells sampled from the same brain regions. Specifically, we integrated our salamander dataset with data from the telencephalon of the agamid lizard *Pogona vitticeps* (34) and from the pallium of the red-eared slider turtle *Trachemys scripta* (which also includes cells from the neighboring subpallium (7)). Clustering of the integrated data yielded 63 clusters, which we refer to as integrated clusters. Hierarchical clustering of average gene expression of integrated clusters produced a cross-species taxonomy of telencephalic neuron types (Fig. 4A-C; Fig. S8).

**Figure 4:**
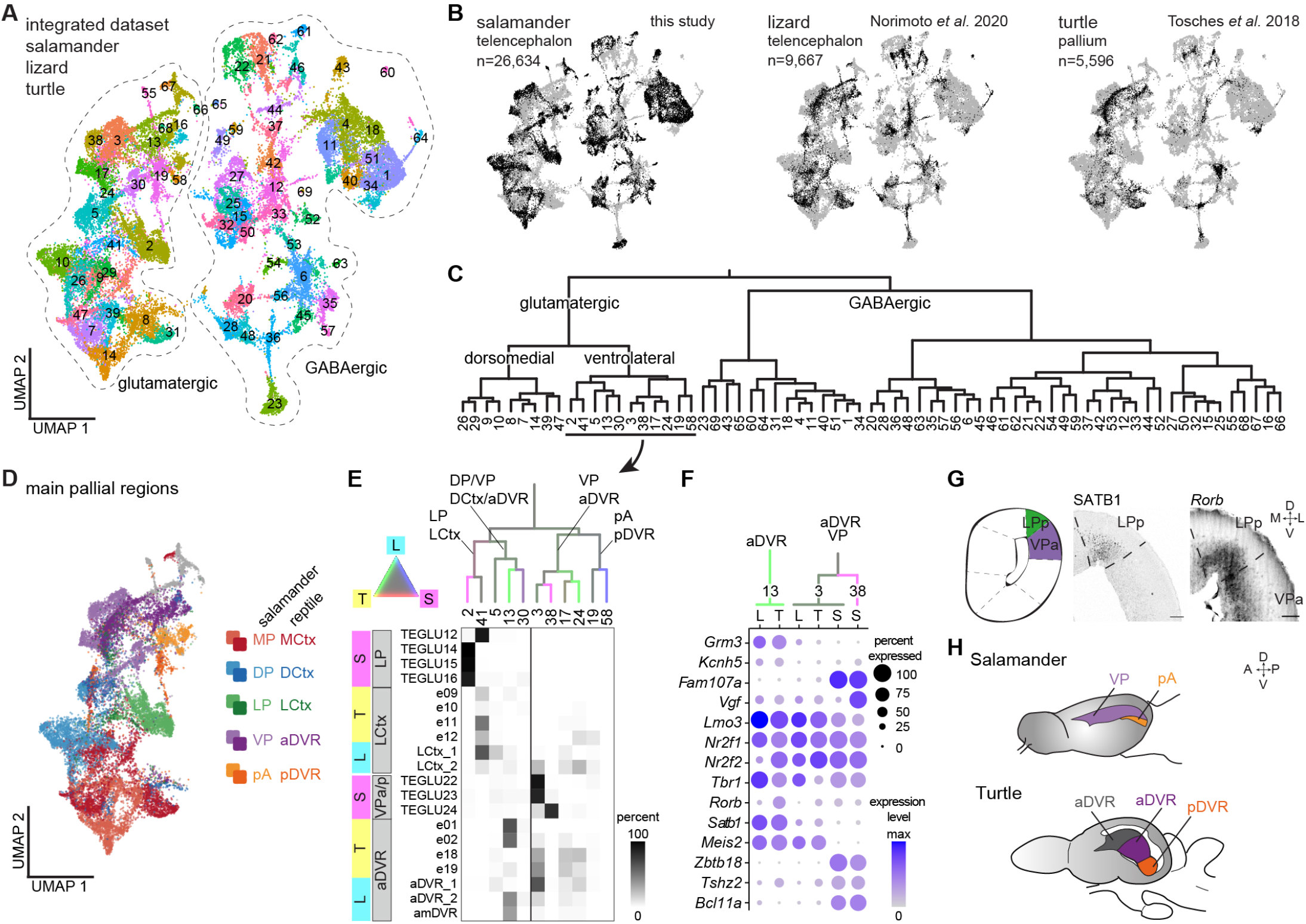
Cross-species comparison of salamander, turtle, and lizard telencephalic neuron types. **(A)** UMAP plot of the integrated scRNAseq dataset from the salamander and lizard telencephalon, and from the turtle pallium, colors indicate clusters in the integrated space. **(B)** UMAP plots of the integrated dataset showing cells from each species highlighted in black. **(C)** Dendrogram showing the molecular similarity of integrated clusters. **(D)** UMAP plot of the glutamatergic clusters from the integrated dataset, colored by pallial region. **(E)** Top: ventrolateral portion of the dendrogram in C, with branches colored by species mixture (gray represents equal proportion of cells from each species). Bottom: percentage of cells from the original species-specific clusters (rows) in the integrated clusters (columns). **(F)** Dotplot showing the expression of molecular markers in aDVR or VP in the integrated clusters, with cells from each integrated cluster split by species (L: lizard, T: turtle, S: salamander). **(G)** Left: schematic of a coronal section at mid-telencephalic level. Right: expression of SATB1 and Rorb in the salamander lateral and ventral pallium. Scale bars: 100um. **(H)** Schematic representation of the salamander and turtle pallium, highlighting ventrolateral domains.

This kind of analysis is built on molecular similarities that result from either homology or convergent evolution. Here, we observed co-clustering of salamander and reptilian cells from pallial regions that are considered homologous, on the basis of independent criteria such as their relative position in the pallium (2, 17). For example, we find co-clustering of salamander MP and the reptilian medial cortex, and of salamander LP and the reptilian lateral cortex (Fig. 4D). Salamander pA and reptilian posterior DVR (pDVR), putative homologs of the mammalian pallial amygdala, co-clustered, while reptilian anterior DVR (aDVR) cells segregated into two integrated clusters. Integrated cluster 3 included salamander cells from the VP, and turtle and lizard cells from the centromedial aDVR (Fig. 4E) (7). Cells in cluster 3 shared expression of several transcription factors, including Tbr1, *Nr2f2, Nr2f1*, and *Lmo3* (Fig. 4F). Integrated cluster 13 comprised only turtle and lizard cells, including cells from the visuo-recipient region of aDVR (7, 35). These reptilian-specific aDVR clusters expressed the transcription factors *Rorb* and *Satb1* at high levels (7), and specific effector genes such as the glutamate receptor *Grm3* (Fig. 4E,F). In *Pleurodeles, Rorb* is expressed at low levels in both LP and VP, and *Satb1* is expressed in LP, but not in DP (Fig. 4G). Taken together, these results indicate that the salamander ventrolateral pallium comprises neuron types with molecular similarity to reptilian lateral cortex and parts of aDVR. Furthermore, they suggest that the sensory-recipient parts of the reptilian aDVR have a unique molecular signature, potentially a result of innovation (Fig. 4H).

### Molecular similarity of salamander dorsal pallium and mammalian entorhinal cortex and subiculum

To extend our molecular comparisons to mammals, we computed gene expression correlations between each salamander telencephalic cluster with digitized *in situ* hybridization data from the Allen Brain Atlas (36) (Fig. 5A). This confirmed the molecular similarity of salamander subpallial regions with their mouse counterparts. Results for the pallium were more ambiguous. For example, salamander LP clusters correlated with hippocampus, neocortex, piriform cortex, and lateral amygdala, pA clusters with the entire pallial amygdala and with piriform cortex, and VP clusters with piriform cortex and lateral amygdala (Fig. 5A). This is consistent with the observation that VP expresses transcription factors with specific or enriched expression in the mouse piriform cortex, such as *Znf536* and *Tshz2* (Fig. 2B, Fig. S6) (36, 37). Using an alternative approach, where we mapped scRNAseq data from the mouse telencephalon (38, 39) on our salamander single-cell dataset (see Methods), we also found correspondences between subpallial regions and hippocampi between the two species. However, using this method, mouse cortical pyramidal types could not be mapped to single salamander clusters, suggesting a high degree of transcriptional divergence (Fig. S9).

**Figure 5:**
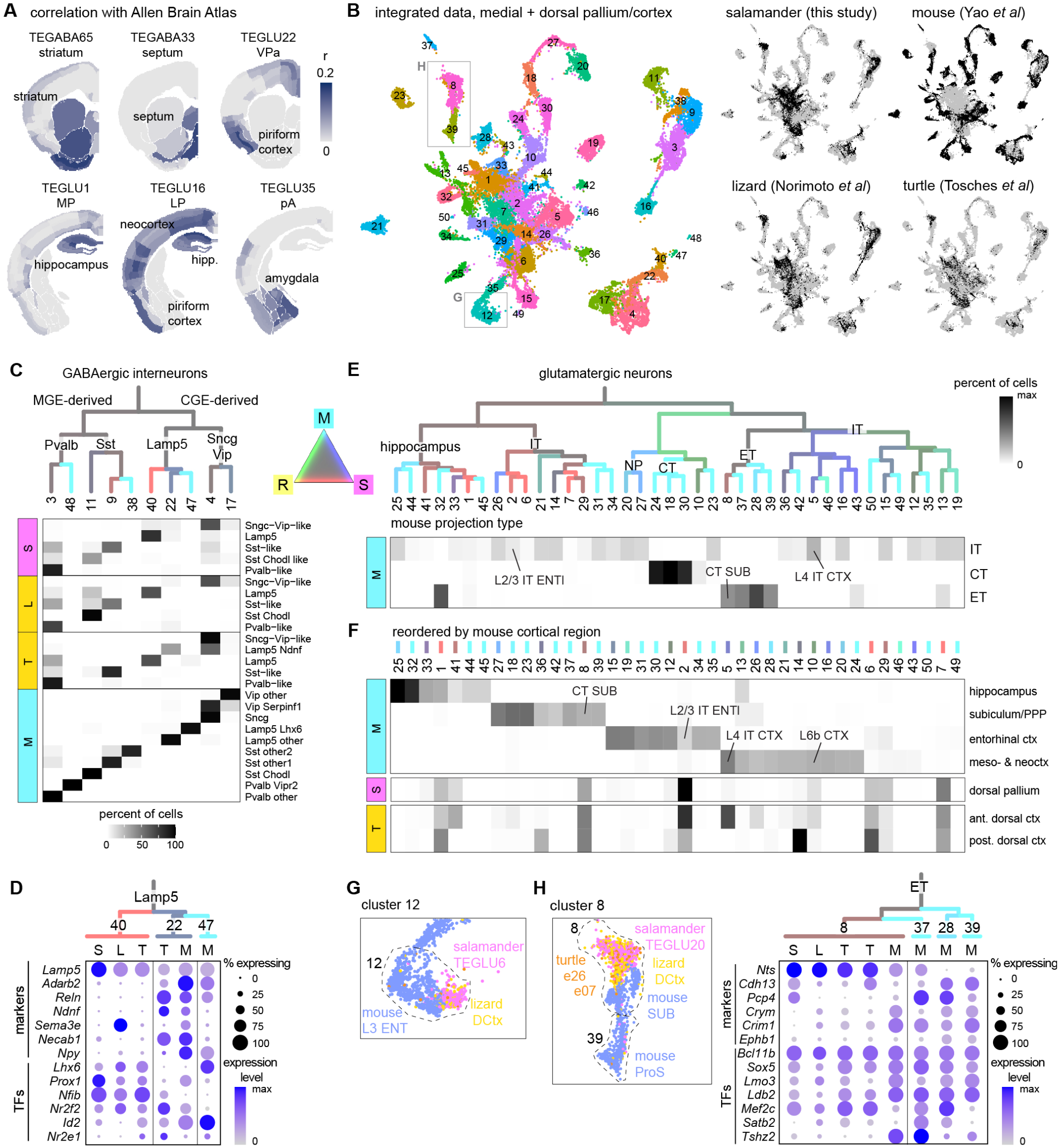
Cross-species comparison of neuron types in the salamander medial and dorsal pallium, reptilian medial and dorsal cortex, and mouse hippocampus and cortex. **(A)** Correlations of the transcriptome of selected Pleurodeles clusters with regional gene expression (in situ hybridization data from the Allen Brain Atlas) in the mouse brain. **(B)** Integration of scRNAseq data from the salamander medial and dorsal pallium, the turtle and lizard (“reptile”) medial and dorsal cortex, and the mouse hippocampus and cortex. Left: UMAP plot of the integrated data, with cells colored by cluster membership. Right: UMAP plots of the integrated dataset showing cells from each species highlighted in black. **(C)** Top: Dendrogram showing molecular similarity of integrated GABAergic clusters, with branches colored by species mixture (gray represents equal proportion of cells from each species). Bottom: percentage of cells from the original species-specific clusters (rows) in the integrated clusters (columns). **(D)** Dotplot showing expression of differentiation markers and of transcription factors (TFs) in Lamp5 INs (integrated clusters 40, 22, and 47). Cells from each integrated cluster split by species (L: lizard, T: turtle, S: salamander, M: mouse). **(E)** Top: Dendrogram showing molecular similarity of integrated glutamatergic clusters, with branches colored by species mixture. Bottom: percentage of mouse cells in each integrated cluster (columns); mouse cells grouped by projection identity (rows). Integrated clusters including selected mouse neuron types are highlighted (L4 IT CTX: thalamorecipient L4 neurons, SUB: subiculum, ENT: entorhinal cortex). **(F)** Top: percentage of mouse cells in each integrated cluster (columns); mouse cells grouped by cortical area (rows), columns reordered by cortical area. Bottom: percentage of salamander DP cells and turtle dorsal cortex cells (rows) in each integrated cluster (columns). **(G)** Close-up on part of the UMAP plot in (B), showing cells in cluster 12 colored by species. **(H)** Left: close-up on part of the UMAP plot in (B), showing cells in cluster 8 colored by species. Right: dotplot showing expression of differentiation markers and transcription factors (TFs) in integrated clusters 8, 37, 28, and 39, split by species.

To identify high-level similarities between salamander, reptilian, and mouse neuron types, we integrated scRNAseq data from derivatives of the dorsomedial pallium, for which complete mouse data are available (39); telencephalic interneurons (INs) (40) were also included in this analysis (Fig. 5B). Salamander GABAergic INs co-clustered with amniote MGE-derived (Pvalb and Sst) and CGE-derived (Lamp5, Sncg, and Vip) IN classes (Fig. 5C; Fig. S9B), indicating that these IN classes trace back to tetrapod ancestors. At deeper levels of IN classification, we found IN types conserved in tetrapods, such as long-range projecting Sst Chodl neurons (cluster 11), and mammalian-specific types, such as Pvalb Vipr2 Chandelier cells (cluster 48) and a subset of Sst INs (cluster 38) (41) (Fig. 5C). Lamp5 INs included a non-mammalian subclass (cluster 40), a turtle and mouse subclass (cluster 22, Lamp5 Ndnf neurogliaform cells in mouse) and a mouse-specific subclass (cluster 47, Lamp5 Lhx6 cells) (41). The transcription factors that differentiate between mammalian Lamp5 Ndnf and Lamp5 Lhx6 INs are coexpressed in non-mammalian Lamp5 cells (cluster 40), suggesting that amniote or mammalian-specific Lamp5 types evolved by diversification of ancestral Lamp5 INs (Fig. 5D).

In the cross-species taxonomy of glutamatergic neurons, major splits corresponded to neurons with distinct projection identities in mouse (IT: intratelencephalic; ET: extratelencephalic, including L5 pyramidal tract neurons; CT: corticothalamic neurons) (Fig. 5E; Fig. S10). Several integrated clusters included only mouse neurons, indicating a greater diversity of mouse pyramidal types. Among these, we found neocortical CT and L5 pyramidal tract neurons, indicating that these neuron types are mammalian innovations (42). In Figure 5F, we plot the same integrated clusters reordered by mouse cortical region, and the proportions of salamander DP and turtle dorsal cortex neurons in these clusters. The large majority of salamander MP and DP neurons co-clustered with mammalian neurons from the hippocampus, entorhinal cortex, and subiculum (Fig. 5F-G, Fig. S10). In contrast, reptilian dorsal cortex neurons also co-clustered with mammalian neocortex, including thalamorecipient L4 IT neurons (Fig. 5F). Consistently, transcription factors instructing neuronal identity in the reptilian dorsal cortex and mammalian neocortex, such as *Satb2* and *Rorb* (43, 44), are not expressed in the salamander DP (Fig. 2B). This indicates that the salamander DP lacks cellular and molecular characteristics of the mammalian neocortex, and is more similar to mammalian cortical areas intercalated between neocortex and hippocampus (13).

### Connectivity of the salamander VP, and similarities with the reptilian aDVR

The results of our comparative analysis prompted us to ask whether neuron types with similar transcriptomes have similar connectivity across species. Expanding on previous findings in salamanders (reviewed in (25)), we conducted retrograde tracing experiments in adult *Pleurodeles* (Fig. 6A). We confirmed that all VP regions project to the putative ventromedial hypothalamus (VMH) homolog (45), and receive afferents from mOB and DP (Fig. 6A-D; Figs. S11,12). Interestingly, VP also receives projections from the central thalamus (Fig. 6A-C), a region expressing *Slc17a6* and *Calb2*, which relays multimodal inputs to the telencephalon and is the amphibian homolog of amniote first-order sensory nuclei (46). These patterns of connectivity resemble the aDVR. In reptiles, aDVR receives inputs from the dorsal cortex, and is organized in subregions innervated by thalamic visual, auditory, and somatosensory nuclei (4, 47–49). Projections to aDVR also include the striatum, the pDVR, and the VMH (4, 47, 50).

**Figure 6:**
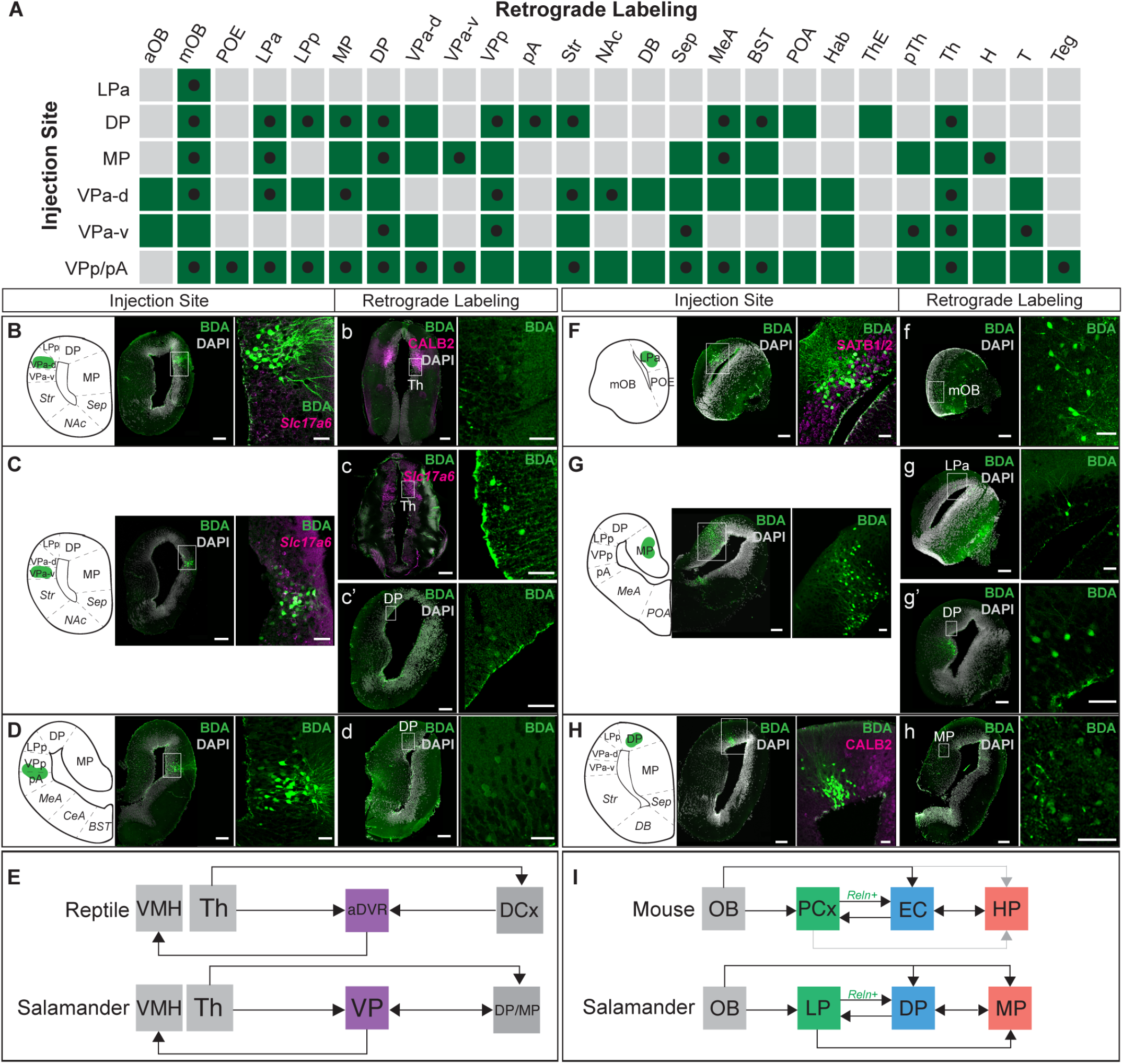
Connectivity of the salamander pallium. **(A)** Summary of regions with retrogradely labeled cells (columns) following 3 kD BDA tracer application into pallial injection sites (rows). Black dots: labeling sites observed in more than one injected brain. **(B-D; F-H)** Left: BDA injection sites, scale bars represent 200 um. Right, magnification of injection site with immunostaining or HCR staining of relevant molecular markers, scale bars represent 50 um. **(b-d, f-h)** left, representative coronal sections where retrogradely labeled cells were identified, with immunostaining or HCR staining of relevant molecular markers when applicable. Right, magnification of retrogradely labeled cells. **(E**,**I)** Schematic connectivity maps of salamander pallial regions depicted above, as compared to mouse (60) and reptile. Gray lines in I represent sparsely documented connectivity. For full list of abbreviations, see Fig. S1; aDVR, anterior dorsal ventricular ridge; BDA, biotinylated dextran amine; DCx, dorsal cortex; EC, entorhinal cortex; HP, hippocampus; mOB, main olfactory bulb; PCx, piriform cortex; Th, thalamus; VMH, ventromedial hypothalamus.

In addition, caudal injections targeting both the salamander VPp (*Slc17a6*+) and pA (*Slc17a6*+, *Lhx9*+) showed more widespread intratelecephalic afferents compared to VPa (Fig. 6A,D; Fig. S11E). Similarly, the reptilian pDVR receives olfactory input and has widespread connectivity with medial and dorsal cortex (48). Thus, the connectivity of salamander VPa and VPp/pA share broad similarities with the reptilian aDVR and pDVR, respectively. While sensory inputs are processed separately by modality in aDVR, there is no indication that this is the case in salamanders (Fig. 6E).

### Connectivity of the salamander MP and DP, and similarities with mammalian hippocampal-entorhinal circuits

In mammals, the primary input to the hippocampal formation is the entorhinal cortex (EC), which includes a lateral EC strongly connected to olfactory areas, and a medial EC, dedicated to processing spatial information (51). The subiculum is the primary output region of the hippocampus. Motivated by our molecular data (Fig. 5), we asked whether the connections between salamander MP and DP are broadly analogous to the connections of mammalian hippocampus, entorhinal cortex, and subiculum, as suggested by previous literature (9, 22, 25). Retrograde tracer injections confirmed that DP and MP are reciprocally connected (Fig. 6A, G,H). MP and DP also receive direct projections from the *Satb1*+/*Reln*+ LP region, an area that receives strong mOB inputs–the lateral olfactory tract runs along this region (23) (Fig. 6F,G; Fig. S12B). Thus, LP neurons are similar to mammalian semilunar cells in the piriform cortex and fan cells in the entorhinal cortex (layer 2), both for their molecular profile (*Satb1, Reln, Lhx2, Tbr1*; Fig. 2 and (21)) and their connectivity (inputs from the mOB, projections to entorhinal deeper layers). Together, these findings suggest that components of mammalian olfactory-entorhinal-hippocampal circuits trace back to tetrapod ancestors (Fig. 6I).

## Discussion

Our molecular data show that, despite its anatomical simplicity, the salamander telencephalon harbors a complex repertoire of neuron types. The combined analysis of their molecular identity, development, and connectivity clarifies the evolution of two novelties in amniotes: the sauropsid aDVR and the mammalian neocortex.

Our data suggest that the aDVR of reptiles and birds comprises at least two sets of neuron types with distinct evolutionary origins. Part of the aDVR traces back to a specialized region of the anterior ventral pallium in the tetrapod ancestor. We propose that this region developed from a ventrolateral pallium field, a progenitor domain distinct from the rest of the pallium. Homologous reptilian aDVR and salamander VPa neuron types express similar gene sets, including a ventral-pallium specific set of transcription factors. A second group of reptilian aDVR neurons does not have an obvious counterpart in *Pleurodeles*; these neurons receive thalamic inputs segregated by sensory modality (visual, somatosensory, auditory, but not olfactory) (35). We propose that neurons in the aDVR specialized in sensory processing are an evolutionary innovation in the sauropsid lineage.

The homologs of these anterior ventral pallium neurons in mammals remain ambiguous. Connectivity data point to similarities of the salamander VPa, aDVR and the mammalian lateral amygdala, a region that receives sensory inputs relayed by the thalamus (47), and expresses some marker genes found in VPa and aDVR, such as *Rorb* (7, 25, 52). However, molecular data indicate similarities of the salamander VPa, bird HVC (part of aDVR (8)), and mammalian piriform cortex (21) (Fig. 5A). A kinship of VPa and aDVR with parts of the mammalian piriform cortex is not entirely surprising, given that the aDVR and the sauropsid olfactory cortex develop sequentially from the same embryonic progenitors (53). Further molecular studies on the mammalian piriform cortex and pallial amygdala, whose cellular diversity remains poorly explored, are needed to clarify their evolutionary relationships with aDVR and VPa.

Our data also clarify the nature of the amphibian dorsal pallium. This region is molecularly distinct from the medial pallium, but does not express many of the markers that define the reptilian dorsal cortex, the area typically compared to the mammalian neocortex for its position, molecular makeup, and connectivity. Our cross-species analysis shows that salamander dorsal pallium neurons cocluster with neurons of the mammalian subiculum and entorhinal cortex. The input-output connectivity of the salamander dorsal pallium suggests that these molecular similarities may correspond in part to conserved circuit motifs. The Reln-expressing neurons in the lateral pallium occupy a peculiar position in this circuit, analogous to Reln neurons in the reptilian olfactory cortex and mammalian piriform (semilunar cells) and entorhinal cortex (fan cells) (21, 54). We propose that mammalian piriform and entorhinal *Reln*-expressing cells are serial homologs (as sister cell types (55)), with the implication that neuron types in layer 2 and in deeper layers of entorhinal cortex may have two distinct evolutionary origins (from the lateral and the dorsal pallium of a tetrapod ancestor, respectively) (20, 56).

In sum, our findings chart the series of innovations that resulted in the emergence of a six-layered neocortex in mammals (Fig. 5). We propose that neocortical L4 Rorb-expressing neurons receiving sensory inputs from the thalamus evolved first, either in amniote ancestors (if salamanders retained the tetrapod ancestral state) (26, 57) or in earlier vertebrate ancestors (with secondary loss in salamanders or amphibians) (58). Neocortical corticothalamic (L6) and pyramidal tract neurons (L5B) emerged later, in mammalian ancestors (42). Expansion of the neocortex led to functional innovations, such as the transition from distributed to columnar information processing (59) and the direct top-down control of locomotion (42). How these molecular and cellular novelties supported such functional innovations within sensory-associative pallial regions remains to be explored.

## Acknowledgements

The authors are grateful to J. Barber, S. Cook, and the Columbia University Institute of Comparative Medicine for animal care; B. Jekely for help with cloning; E. Subramanian for contributing to the IsoSeq reference; A. Matheson for the diagram in Fig. 6I, and A. Matheson, L. Xu, G. Laurent, O. Hobert, and R. Satija for critical feedback on the manuscript. Light-sheet imaging was performed with support from L. Hammond and the Zuckerman Institute’s Cellular Imaging platform (NIH 1S10OD023587-01); computing resources were provided by Columbia University’s Shared Research Computing Facility (NIH 1G20RR030893-01 and NYSTAR Contract C090171). The authors acknowledge support of the National Genomics Infrastructure (NGI)/Uppsala Genome Center and UPPMAX for providing assistance in massive parallel sequencing and computational infrastructure. Work performed at NGI / Uppsala Genome Center has been funded by RFI/VR and Science for Life Laboratory, Sweden.

## Author contributions

scRNAseq data: JW, AOG, AD, EG; scRNAseq analysis: EG, AOG, MAT, JG; Iso-Seq transcriptome: AJA, NL, AS, MY; anatomy and histology: JW, AD, ECJ, GM; axonal tracing: JW, ECJ; manuscript writing: MAT, AD, ECJ, AOG, JW, EG, with edits from AJA; project management and supervision: MAT.

## Materials and Methods

### Animals

Adult *Pleurodeles waltl* were obtained from breeding colonies established at Columbia University and Karolinska Institute. Animals were maintained in an aquatics facility at 20°C under a 12L:12D cycle (61). All experiments were conducted in accordance with the NIH guidelines and with the approval of the Columbia University Institutional Animal Care and Use Committee (IACUC protocol AC-AABF2564). Experiments were performed with adult (5-19 months) male and female salamanders, and stage 36, stage 41 and stage 50 embryos and larvae (staged according to the Gallien and Durocher 1957 atlas (62)).

### Brain dissociation and single-cell capture

For adult brain dissociation, animals were deeply anesthetized by submersion in 0.2% MS-222. The brain was perfused transcardially with ice-cold oxygenated Amphibian Ringer’s solution (96 mM NaCl; 20 mM NaHCO3; 2 mM KCl; 10 mM HEPES; 11 mM glucose; 2 mM CaCl2; 0.5 mM MgCl2), and the animal decapitated. The whole brain or telencephalon was dissected out, and embedded in 4% LM Agarose in Ringer. The brain was then sliced coronally on a vibratome into 500 um sections, and cut into 500 um cubes in cold carbogenated Ringer. The tissue pieces were incubated in dissociation buffer (Papain, DNase, Liberase, TTX), and then mechanically dissociated with silanized glass pipettes of decreasing tip diameter. The supernatant was then strained with carbogenated calcium-free Hibernate A media (BrainBits). To filter out cell debris, the cell suspension was centrifuged through a BSA density gradient, supernatant removed, and then resuspended. The cells were then diluted (approx. 1000 cells/µL) in calcium and magnesium-free Hibernate A. Cells were loaded into a 10x Chromium Chip G with a targeted cell recovery of 5000-8000 cells for GEM Generation and cell barcoding. Single-cell RNA-seq libraries were prepared with 10x Chromium Next GEM Single Cell 3ʹ Reagent Kit v3.0 or v3.1 (Dual Index). For the developmental scRNA-seq dataset, telencephali from 35 larvae at stage 36, 10 larvae at stage 41, and 5 larvae at stage 50 were isolated, and the Tissue pieces were dissociated for 30 min at room temperature using the Papain dissociation kit (Worthington) in calcium-free Hibernate A (BrainBits). After processing and concentration, cells were resuspended in calcium and magnesium-free Hibernate A (BrainBits) at a concentration of 1000 cells/µl. Cells were counted in a haemocytometer chamber and immediately processed for single-cell GEM formation (10x Genomics, single cell RNA sequencing 3’, Chromium V3.1). Cells from stage 36 larvae were multiplexed using 4 tags from the Chromium Next GEM Single Cell 3ʹ v3.1 prior to GEM formation.

### Reference transcriptome

Tissue was harvested from the brain of a female adult *P. waltl*. Tissue was immediately frozen on dry ice and stored at -80°C. Approximately 25 mg of brain tissue was then used for tissue pulverization in liquid nitrogen. Tissue powder was used for RNA extraction using Total RNA Purification Kit (Cat. 17200 from Norgen Biotek) as per manufacturer’s recommendation. Genomic DNA was removed via on column DNA removal using Norgen RNAse free DNAase kit (Cat. 25710). RNA quality was analyzed via bioanalyzer and had a RIN value of 10. For IsoSeq, cDNA was synthesized using SMARTer PCR cDNA (Clonetech, CA). One IsoSeq library was prepared. Sequencing was performed on a single SMRT cell using PacBio Sequel II. This produced 3,946,27 reads with a mean number of passes of 19, and average read length of 3,618 bp. Data was then fed into IsoSeq3 (https://github.com/PacificBiosciences/IsoSeq) to generate fasta outputs of high quality and low quality reads. This generated 287,253 polished high-quality isoforms, and 2,802 polished low-quality isoforms. The high quality fasta output was taken for further use as a reference transcriptome. This IsoSeq transcriptome and previously published short-read based de novo transcriptome (63), were used to generate a combined reference transcriptome. TransDecoder was used to generate predicted peptide sequences for each IsoSeq transcript (64). For transcripts with multiple predicted peptide sequences, only the sequence with the lowest e value was kept. These predicted peptides were provided to EggNOG-mapper (v2.16), along with the published iNewt peptide sequence to generate a transcript to gene name mapping (65). All parameters were set to default (Taxonomic Scope = Auto, Orthology Restrictions = Transfer annotations from any orthologue, Gene Ontology Evidence= Transfer non-electronic annotations, PFAM refinement = Report PFAM Domains from Orthologues, SMART annotation = Skip). Peptide sequences that did not receive a “preferred name” from the EggNOG-mapper were removed. This would occur when either the sequence was not assigned an orthologous sequence, or the assigned sequence was not characterized enough to receive a common gene name. The name for the RNA transcript sequence that correlated with the peptide sequence used for EggNOG mapping was matched with the corresponding “preferred name” from the eggnog output to generate a tgMAP for subsequent Alevin analysis (described below) (66). The merged transcriptome was then filtered to only contain genes which were included in this tgMAP file (again, based on their assignment of a “prefered name” from EggNOG). This final merged reference transcriptome contained 194,403 transcripts of 17,111 uniquely annotated genes.

### Single-cell read alignment

Developmental Bioinformatic Processing: Reads were assigned to the combined *Pleurodeles waltl* transcriptome generated above using Alevin (Salmon v1.6.0) (66). Library type was set to automatic and keepCBFraction was set to 1. All other parameters were set as default. The stage 36 library was multiplexed (as described above) and required demultiplexing prior to analysis. For this, multiplex tag reads were also assigned using Alevin.

### Single-cell data QC and filtering

For adults: Count matrices were obtained after alignment with Alevin (Salmon v1.6.0) and knee plots were generated for each library. Each library was processed independently, filtering out droplets with low UMI counts and individual Seurat objects were generated per library using Seurat 4.0.2. After filtering, all objects were merged together into a single Seurat object containing all libraries. We calculated the percentage of all counts that belonged to mitochondrial genes and filtered our dataset to keep the cells that had a mitochondrial gene expression below 15% and more than 800 genes per cell, obtaining a dataset of 52,638 cells. For further filtering of low-quality cells, the dataset was screened for low-quality cell clusters (low number of genes per cell detected and high percentage of mitochondrial gene expression). Additional screening of bad clusters was performed on the basis of their marker genes, annotating a cluster as low-quality whenever it did not express any cluster-specific marker gene (these low-quality clusters expressed high levels of housekeeping, mitochondrial, and ribosomal genes). Subsequently, additional bad-quality cells were identified using a Support Vector Machine (SVM) classifier (7). The SVM was trained using ∼10% of the cells from all clusters identified as low-quality clusters, and the same number of cells from the good clusters. This allowed the SVM to find low-quality cells in the entire dataset and remove them, generating a final Seurat object of 36,116 cells. Additional quality control of this final dataset was performed by analyzing the number of genes per cell, number of UMIs per cell, and percentage of mitochondrial gene expression both by library source and by animal source. For Development: The same protocol was used as for the adult brain, except that cells from stage 41 and 50 telencephalon samples were initially filtered at 1000 reads and 15% reads from mitochondrial genes, before being cleaned by SVM as described above. Cells from the stage 36 telencephalon were initially filtered at 700 reads before demultiplexing using the Seurat package. Only cells labeled as singlets were kept for further analysis, and were cleaned by SVM before integrating with the stage 41 and 50 samples.

### Single-cell clustering

The adult 36,116-cell dataset was clustered and analyzed using Seurat 4.0.2 (67). Raw data was normalized using the SCTransform function from Seurat, regressing the data by the following variables: animal, percentage of mitochondrial gene expression, number of genes per cell and number of UMIs per cell. PCA was calculated using 200 PCs. Based on the Elbow Plot from PCA, we proceeded downstream with the analysis using 50 dimensions for calculating our UMAP. We used 2000 variable genes for finding neighbors and finding clusters. For clustering, we used the default clustering method and the original Louvain algorithm with res = 2. Using well-known marker genes, the original clusters were merged into 11 clusters, corresponding to major neuronal and non-neuronal cell classes. Subsequently, this dataset was subsetted to contain only neurons, identified as clusters expressing neuronal markers such as *Syt1, Snap25, Slc17a7, Slc17a6, Gad1* and *Gad2*. This neuronal dataset was processed as described above, normalizing the data with the SCTransform function from Seurat, regressing out animal, percentage of mitochondrial gene expression, number of UMIs per cell and number of genes per cell and calculating PCA with 350 PCs. Similarly, based on the Elbow Plot we proceeded with the downstream analysis using 180 dimensions. We used 2000 variable genes, Louvain algorithm and resolution = 6 for calculating neighbors and finding clusters. At this resolution, parts of the dataset were overclustered, as indicated by the fact that adjacent clusters on the UMAP space corresponded to cells sampled from different animals. These clusters were manually curated and merged if they did show differential expression of marker genes. After these curations steps, the final clusters were renamed according to their excitatory (*Slc17a7, Slc17a6*) or inhibitory (*Gad1, Gad2*) identity, as well as their anatomical location (telencephalon or diencephalon): TEGLU refers to telencephalic glutamatergic neurons, TEGABA to telencephalic GABAergic neurons, and DIME to diencephalic and mesencephalic neurons. This final neuronal dataset contained 29,294 cells and 114 clusters. Developmental data: SVM cleaned data from stage 36, 41 and 50 was combined into one Seurat object using the “merge” command, which contained 8,615 cells over 30 clusters. Non-neuronal clusters were then removed on the basis of immunological and hematopoietic marker genes. Cell cycle scores were computed for the remaining 8,510 neurons, using the CellCycleScoring command based on Seurat’s built in list of cell cycle genes. These neurons were normalized using the SCTransform command based on 3000 genes, while regressing for RNA count, stage, percent mitochondrial genes, G1 score, and G2M score. Clusters were calculated based on 25 dimensions, with a clustering resolution of 2 to generate 29 clusters. This sample was further cleaned for trajectory inference by removing non-telencephalic clusters based on lack of expression of the telencephalic cluster *Foxg1* (68). Label transfer against the adult data: Label transfer within the Seurat package was performed to annotate the developmental clusters based on differentiated adult cell types. This feature identifies transfer anchors between two data sets to allow a comparison of cell type identity between the two, meaning that developmental cells can be classified based on which adult cell they are most similar to (as in figure 5D). FindTransferAnchors was run using dims= 1:50 and k.anchor=10. TransferData was run using k.weight=20. The remaining 7782 telencephalic cells were clustered as above. To generate Figure S10F, the MapQuery function was used to plot the developmental cells on the adult UMAP space. For plotting, a prediction score threshold of 0.9 was used.

### Hierarchical clustering of adult neuron types

An average expression matrix was calculated for the neuronal object using the SCTransform normalized data, keeping only the top 2000 variable genes. Correlation was calculated using the Spearman correlation method and hierarchical clustering was performed using Ward’s minimum variance method, squaring dissimilarities before clustering (‘ward.D2’).

### Trajectory inference

The developmental telencephalic clusters were annotated according to their corresponding pallial region as described above. *Neurog2* expression was used to identify pallial progenitors, as this gene is expressed predominantly in the ventricular zone, mediating pyramidal neuron specification in vertebrates (69, 70). Clusters that corresponded to DP, MP, LP, VP, and MT, as well as their progenitors, were subsetted without reclustering. For this trajectory analysis, we excluded cells from the pA, since we limited our dissection of the telencephalon to the evaginated telencephalic vesicles. Therefore, we did not sample pA progenitors located in the caudal portion or in the non-evaginated part of the telencephalon, and trajectories cannot be calculated. The subset was provided to Slingshot (v1.8.0) (31) for trajectory analysis using default parameters, and pseudotime was calculated along each of the inferred trajectories by measuring the distance of each cell along a Slingshot-defined trajectory from a common progenitor population to one of four terminal branches. Genes that define the dorsomedial and ventrolateral trajectories were determined by comparing clusters along each trajectory with a corresponding cluster in the other, using the FindMarkers function within Seurat (default parameters-gene.use set to all, and test.use set to bimod). Genes were selected for heatmap plots in Fig. 5 if they showed greater than 21% difference in the absolute proportion of cells expressing these genes between clusters, and 3 fold greater percentage of cells that express the gene between the clusters that compose each trajectory. p values for all genes identified were below 1e-9. Genes were then ordered on the heatmap based on when they reached peak expression along pseudotime. A heatmap was generated using the ComplexHeatmap package (71), and heatmap colors were assigned based on the root mean square of cell expression levels for each gene using the scale function.

### Cross-species comparisons of transcriptomics data

Using Seurat’s integration pipeline, we generated two integrated datasets. The first integrated dataset was generated from lizard (34), turtle (7) and salamander (this study) scRNA-Seq data. For consistent comparisons, we subsetted the original datasets in order to only include cells that were sampled from equivalent brain regions. In this case, we kept all the cells from lizards (Norimoto et al 2020), and excitatory and inhibitory clusters from turtles which included interneurons and some subpallium (Tosches et al 2018), excluding the unidentified clusters. For salamanders, we included cortical pallium, septum, striatum, interneurons and olfactory bulb inhibitory neurons, excluding glutamatergic cells from the olfactory bulb, because the olfactory bulb was not sampled in the reptilian datasets. After subsetting, each dataset was normalized independently using Seurat’s SCTransform function. For each dataset, a different list of variables were provided for regression (lizard: percent of mitochondrial genes, number of genes per cell, number of UMIs per cell; turtle and salamander: percent of mitochondrial genes, number of genes per cell, number of UMIs per cell, animal of origin). These three datasets were brought together in a list class object, from which the integration features were calculated. Using the SelectIntegrationFeature function in Seurat, we selected the top 1000 genes that are variable across datasets and prepared the dataset for integration using these 1000 genes. Cells matched in the three datasets were identified (anchors) by using FindIntegrationAnchors function using Canonical Correlation Analysis (CCA) and normalizing with SCT. Integration was carried out with the Seurat function IntegrateData, normalizing with SCT, using the anchor sets obtained from FindIntegrationAnchors and using 80 dimensions. Downstream processing was performed as described above, calculating PCA with 200 PCs and UMAP with 80 dimensions. Clustering analysis was performed with default method, using SLM algorithm and clustering res = 2.1. The resulting integrated object consists of 42,007 cells from the three species, with a total of 69 clusters from which 20 are glutamatergic clusters and 49 GABAergic clusters. The second integrated dataset included cells from lizard (Norimoto et al 2020, (34)), turtle (Tosches at al 2018, (7)), mouse (Yao et al 2021, (39)) and salamander (this study). In order to make meaningful comparisons, we subsetted the datasets in order to include cells that had a matched cell population, as described above. The target for this integration was cells from the medial pallium and dorsal pallium. For lizards, we subsetted the dataset to keep only medial cortex (MCtx), dorsal cortex (DCtx), MGE-derived interneurons and LGE-derived interneurons. Similarly, the turtle dataset was subsetted to keep cells from the dorsal cortex (DC), medial cortex (MC), dorsomedial cortex (DMC), and interneurons. For salamanders, we included medial pallium, dorsal pallium, ventral pallium, and interneurons. The mouse dataset was subsampled to contain only 25,000 cells from the original dataset, keeping only cells from hippocampus, neocortex/mesocortex, subiculum, entorhinal and cortical interneurons. Following a similar approach to what described above, all datasets were independently normalized using SCT. For turtle, lizard and salamander, the same variables to regress were used. For mouse normalization, variables to regress were number of genes per cell, number of UMIs per cell and external_donor_name_label. To select genes for integration, we used a different approach given that the diversity of mammalian cells biases the integrated features towards mammalian genes. For balancing the list of integration genes, we calculated the top 5000 variable genes from each individual dataset and took the intersection of these lists. Additionally, we removed from the integration list any gene that had a ‘salt-and-pepper’ expression in any dataset using the NoisyGene function and intersected all the lists of noisy genes from each species. The remaining number of genes to carry out the integration was 897 genes. The rest of the analysis was carried out as described above. The final integrated dataset for all four species consisted of 38,103 cells and 50 clusters. To identify molecular similarities among different species’ cells in each integrated cluster, we performed a cross-species taxonomy analysis for both integrated datasets, using the tree-based method and package previously developed by Bakken et al 2021 (https://github.com/huqiwen0313/speciesTree). Similarities of integrated clusters were calculated as 1-cor(x) using the Spearman correlation method and hierarchical clustering of this obtained matrix was performed with Ward’s method. The resulting dendrogram represents the relationship between the different integrated clusters based on gene expression of the cells contained in the integrated cluster, color-coded according to the proportion of each species’ cells in the integrated cluster.

### Tissue preparation for histology on sections

Animals were deeply anesthetized, with adults submerged in 0.2% with MS-222 and larvae submerged in 0.04% MS-222. All solutions during tissue preparation for in situ hybridization were prepared cold in RNase-free solutions prepared with DEPC-treated H2O. Adult animals were transcardially perfused with 10 mL PBS, followed by 10 mL 4% PFA in PBS, then decapitated. Brains were dissected out, and postfixed overnight at 4°C in 4% PFA in PBS. Postfixation was stopped in PTW (PBS with 0.1% Tween). Coronal sections were prepared using a Leica VT1200S vibratome (70 um thickness) with the brain embedded in 4% low-melting agarose. Larvae were fixed overnight at 4°C in 4% PFA in PBS, washed in PBS and cryoprotected in 30% sucrose-PBS. After embedding in Tissue-Tek OCT compound (Sakura), 12-20 um coronal sections were cut on a cryostat and mounted on glass slides.

### Immunohistochemistry on sections

Sections were blocked in Blocking Buffer (2.5% BSA, 2.5% sheep serum, 50 mM glycine) in PBST (PBS with 0.2% Triton), then incubated with mouse anti-NeuN (1:500, Sigma-Aldrich MAB377), mouse anti-SATB2 (1:50, abcam ab51502) and/or rabbit anti-SOX2 (1:500, abcam ab97959) in primary Ab solution (10 mM glycine, 0.1% H2O2 in PBST) 1-3 nights at 4°C. Importantly, our scRNAseq analysis and in situ hybridization indicate that abcam ab51502 mouse anti-SATB2 recognizes salamander SATB1 instead. Specifically, Satb1 in situ hybridization produced staining in the LPa and LPp, consistent with co-expression of *Satb1, Reln* and *Lhx2* in the lateral pallium. Additionally, Satb2 in situ hybridization did not result in any staining, consistent with the fact that Satb2 was not detected in scRNAseq data in pallial neurons. Therefore, in the main text, we will be referring to the SATB2 antibody as SATB1. The tissue was washed 5 × 15 min in PBST at RT, then incubated in goat anti-mouse IgG, goat anti-rabbit IgG conjugated to Alexa 488, Alexa 594, or Alexa 647 (1:500, Invitrogen) with DAPI (1:5000) in PBST overnight at 4 °C or for 2 hrs at room temperature. The tissue was washed 5 × 15 min in PBST at RT. The tissue was washed once more in PBST and mounted in DAKO fluorescent mounting medium (Agilent Technologies). Images were acquired using a confocal microscope (Zeiss LSM800) and processed in FIJI.

### *In situ* hybridization on sections

Vibratome sections were postfixed overnight at 4°C in 4% PFA in PBS, washed in PTW (PBS with 0.1% Tween-20), and the pia was peeled off manually with Dumont 5SF forceps (FST). Floating tissue sections were permeabilized for 8 min in 10 ug/mL ProK, washed 2 × 2 min in 2 mg/mL glycine, and rinsed in PTW for 5 min. To block nonspecific binding, the sections were acetylated by incubation for 5 min in 1% TEA/PTW, followed by 5 min in 1% TEA and 3 uL/mL acetic anhydride in PTW, and a 5 min wash in PTW. This was followed by postfixation for 20 min at RT, and 3 × 10 min washes in PTW. All stock solutions were prepared in DEPC treated or Molecular grade water. To prepare probes for ISH, ∼1kb fragments from coding regions of genes of interest were PCR amplified from a *Pleurodeles waltl* brain cDNA library or ordered from Twist Bioscience. Fragments were cloned into the pCRII vector and sequences verified by Sanger sequencing (Eton Bio). After plasmid linearization, anti-sense DIG-labeled RNA probes were generated by in vitro transcription and purified with the RNeasy kit (Qiagen). At 55-62°C, the tissue was pre-hybridized for 1 hr in Hybridization Mix (50% formamide, 5x SSC, 50 ug/mL heparin, 250 ug/mL yeast tRNA, 5x Denhardt’s solution, 0.2% Tween, 500 ug/mL salmon sperm DNA, 10% Dextran sulfate in DEPC-treated water). The sections were then incubated 1-2 nights in Hybridization Mix with denatured riboprobes (1-3 ng/uL). Following hybridization, the sections were washed 2 × 30 min in a pre-warmed low stringency wash buffer (50% formamide, 2x SSC, 0.1% Tween in DEPC-treated water), 2 × 40 min in prewarmed high stringency wash buffer (0.1 - 0.2x SSC, 0.1% Tween), and 2 × 10 in MABT (100 mM maleic acid, 150 mM NaCl, pH 7.5) at RT. The sections were blocked in Blocking Buffer (5% sheep serum with 10% Roche Blocking Reagent 10x in MABT) for 1 hr at RT and incubated overnight with anti-Digoxigenin-AP (1:4000, Roche 11093274910) in Blocking Buffer. Signal was developed by washing 4 × 30 min in MABT, followed by incubation in fresh staining solution (alkaline phosphatase buffer pH 9.5, 4.5 ug/mL NBT, 3.5 ug/mL BCIP, 5% polyvinyl alcohol) for 1-5 days. The staining reaction was stopped in PBS pH 7.4, and the sections were mounted in DAKO fluorescent mounting medium (Agilent Technologies). Images were acquired using an upright brightfield microscope (Leica DMR with Basler color camera, ACCU-Slide MS software). Background was subtracted evenly across the section using Photoshop CS6, and air bubbles accidentally present outside of the tissue section (for example, in the ventricle) were cropped out.

### Hybridization chain rection (HCR) *in situ* hybridization on sections

HCR-3.0-style probe pairs for fluorescent in situ mRNA detection were ordered from Molecular instruments or designed using the insitu_probe_generator (72) and ordered from IDT (20-33 pairs per probe set). After the same tissue pretreatment described above, the Molecular Instruments HCR v3.0 protocol for sample in solution (rev. 6) was followed (73). At 37°C, the tissue was pre-hybridized for 30 min in probe hybridization buffer (Molecular Instruments), and incubated overnight in probe hybridization buffer with 4-20 pmol of probe sets (*Slc17a6*). Excess probe was removed by 4 × 15 min washes in probe wash buffer (Molecular Instruments) at 37°C, and 3 × 5 min washes in 5x SSCT at RT. The sections were preamplified for 30 min in amplification buffer (Molecular Instruments), and incubated overnight in amplification buffer with snap-cooled hairpins (60 pM, Molecular Instruments) in the dark at RT. Excess hairpin was removed by 2 × 5 min, 2 × 30 min, 1 × 5 min washes in 5x SSCT at RT. Sections were incubated 1 hr in 1:5000 DAPI in 5x SSCT, washed 3 × 5 min in 5x SSCT, and mounted in DAKO fluorescent mounting medium (Agilent Technologies). Images were acquired using a confocal microscope (Zeiss LSM800) and processed in FIJI.

### Hybridization chain rection (HCR) *in situ* hybridization, iDISCO brain clearing, and visualization of wholemount samples

Wholemount tissue staining and clearing was based on iDISCO (see (74) and DIIFCO (HCR, see (73, 75) protocols. Following fixation, the brains were rinsed 3 × 15 min in PBS. For clearing, the tissue was treated with an increasing gradient of methanol in PBS (20%, 40%, 60%, 80%, 100%, 1 hr each), bleached overnight with 5% hydrogen peroxide in 20% DMSO/methanol at 4°C, then rehydrated with a decreasing gradient of methanol. At 37°C, the brains were permeabilized for 1d in 0.2% Triton X-100, 20% DMSO and 0.3M glycine, the tissue was pre hybridized for 30 min in probe hybridization buffer (Molecular Instruments), and incubated 2 nights in probe hybridization buffer with 2-4 pmol of each probe set (*Etv1, Sox6, Slc17a6, Nr2f2, Penk, Rorb*). Excess probe was removed by 3 × 1 hr washes in probe wash buffer (Molecular Instruments) at 37°C, and washed overnight in 5x SSCT at RT. The tissue was preamplified for 30 min in amplification buffer (Molecular Instruments), and incubated 2 nights in amplification buffer with snap-cooled hairpins (60pM, Molecular Instruments) in the dark at RT. Excess hairpin was removed by 3 × 1 hr washes, followed by an overnight wash in 5x SSCT. The samples were embedded in 4% agarose, dehydrated in a decreasing methanol gradient, and incubated in 66% DCM/ 33% methanol for 3 hr at RT. Residual methanol was removed with 2 × 15 min washes in 100% DCM, and the tissue was allowed to clear overnight in DBE. Images were acquired using a LaVision Ultramicroscope II light sheet microscope at 4X magnification and 2 µm resolution, and subsequently visualized using ImarisViewer 9.8.0. Oversaturated impurities on the brain surface or within the ventricle were masked based on fluorescence intensity.

### Axonal tracing

To provide access to all injection sites, and in accordance with similar tract-tracing experiments in amphibians (16), all injections were conducted *ex vivo*, under which conditions *Pleurodeles* brain tissue survives up to 2 days. Prior to *ex vivo* tracer injection, animals were deeply anesthetized by submersion in 0.2% MS-222. The brain was perfused transcardially with ice-cold oxygenated Amphibian Ringer’s solution (96 mM NaCl; 20 mM Na-HCO3; 2 mM KCl; 10 mM HEPES; 11 mM glucose; 2 mM CaCl2; 0.5 mM MgCl2), and the animal decapitated. Brains were subsequently exposed while inside the skull, and the dura mater and choroid plexus removed. The arachnoid and pia mater were then removed using fine forceps only above the brain region to be injected. 10% 3 kD or 10 kD biotinylated dextran amines (3 kD BDA, Invitrogen D7135; 10 kD BDA Invitrogen D1956; diluted in 0.9% NaCl) were used as retrograde and anterograde tracers, respectively. While the two molecular weights do not result in exclusive unidirectional transport, the robust differential labeling of cell bodies (retrograde) versus axons (anterograde) depending on the injected construct was clear during image analysis, and has been used historically (76, 77). All tracer solutions were pressure injected (20-30 nL, 1 nL/sec) using a glass capillary needle (diameter ∼10 µm) connected to a Nanoject III injection system (Drummond); after injection, the needle was left in the injection site for 5 min to prevent leakage. Injection sites (OB (n=2), LPa (n=2), DP (n=2), MP (n=2), VPa-v (n=2), VPa-d (n=2), VPp/pA (n=2), VMH (n=1)) were determined using anatomical landmarks, and successful tracer application was confirmed with FastGreen dye (Sigma-Aldrich F7252). Injected brains were transferred to fresh ice-cold Amphibian Ringer’s solution and allowed to incubate for 24-48 hr on ice, with constant oxygenation. Brains were then fully extracted from the skull, transferred to 4% PFA, and fixed overnight at 4 ºC. After fixation, 70 µm sections were cut on a vibratome, and processed either according to IHC or HCR staining protocol, as described above. During secondary antibody incubation (IHC) or during hairpin amplification (HCR), 1:500 Streptavidin conjugated to Alexa Fluor 488, 594, or 647 (Invitrogen S32354/S32356/S32354) was added to the solution to visualize BDA localization. All slices were subsequently mounted onto glass slides with DAKO fluorescent mounting medium (Agilent Technologies). Images of each section were acquired using a confocal microscope (Zeiss LSM800) and labeling was scored manually by two independent researchers. Localization and annotation of BDA tracer signal was determined using co-staining for known molecular markers, as well as anatomical landmarks (Fig. S1, Movie S1).

## Supplementary Figures

**Fig. S1.**
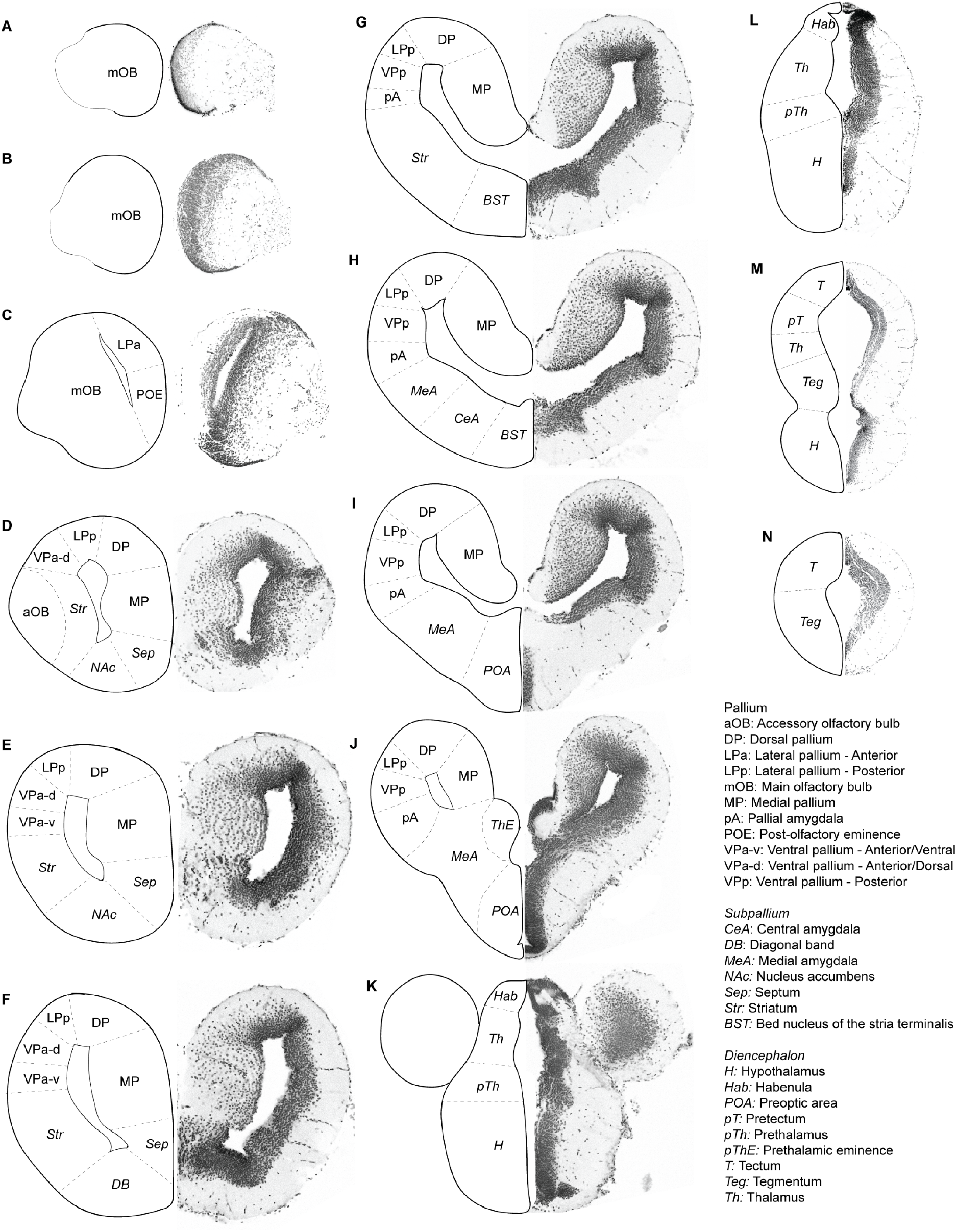
DAPI atlas. Coronal sections (70 µm thick) of the *Pleurodeles waltl* brain, arranged anterior **(A)** to posterior **(N)**. Nuclear (DAPI) stain is shown on the right in each panel, with a schematic showing subdivisions on the left, annotated with the acronyms described in the key. Annotations were performed using histological data from the present study, in tandem with previous literature (9, 10, 78-81). See also Movie S1.

**Fig. S2.**
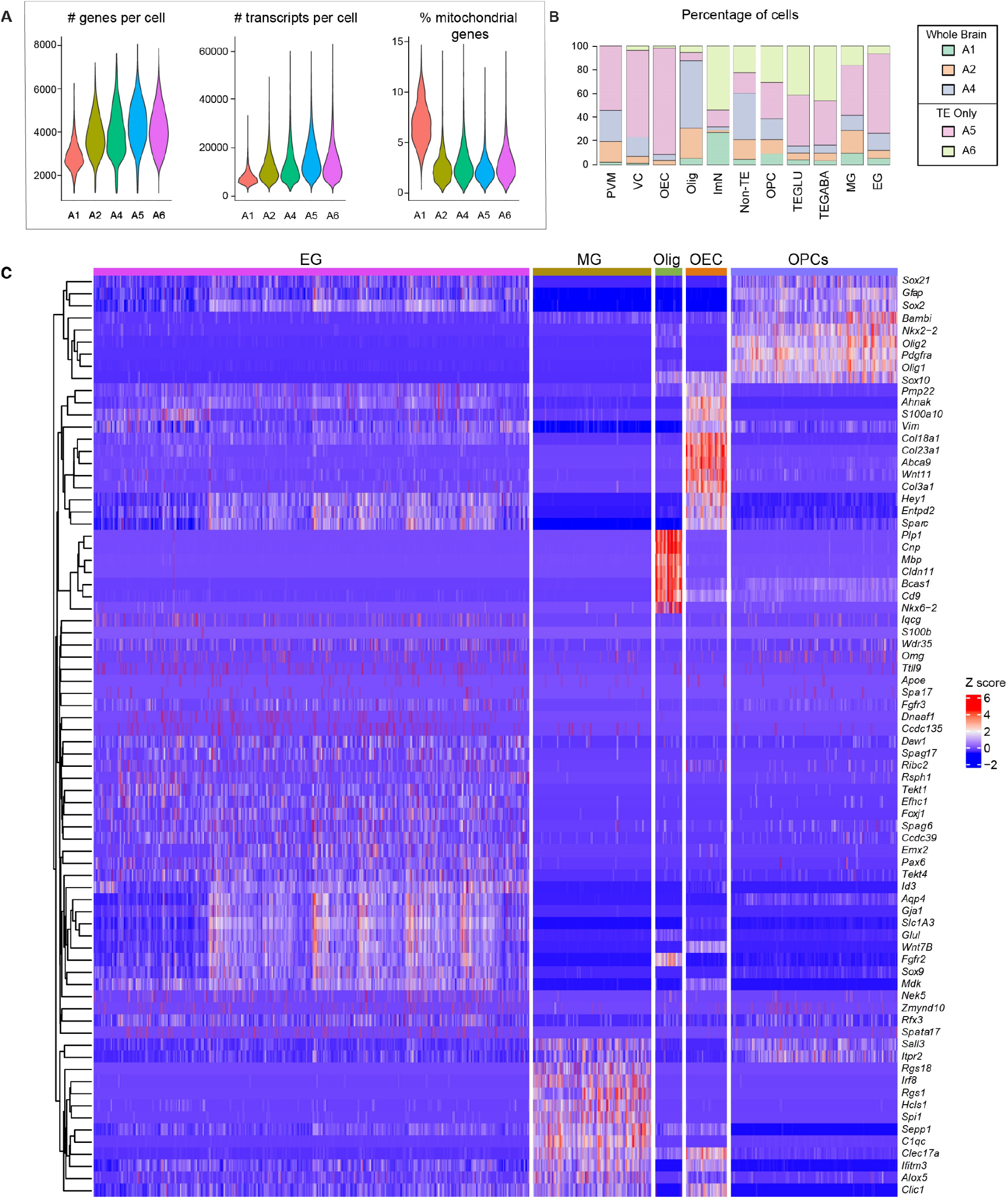
Quality control of the adult brain single-cell dataset and additional gene expression profiles of non-neuronal cells. **(A)** Violin plots of the number of genes or transcripts per cell, and percent mitochondrial genes in the global dataset, sorted by animal. **(B)** Bar plot of percentage of cells per animal in global dataset clusters. Legend denoting tissue included for each animal in single cell prep for dissociation. **(C)** (left) Dendrogram hierarchically clustered based on molecular similarity. (right) Heatmap of differentially expressed genes for non-neuronal clusters.

**Fig. S3.**
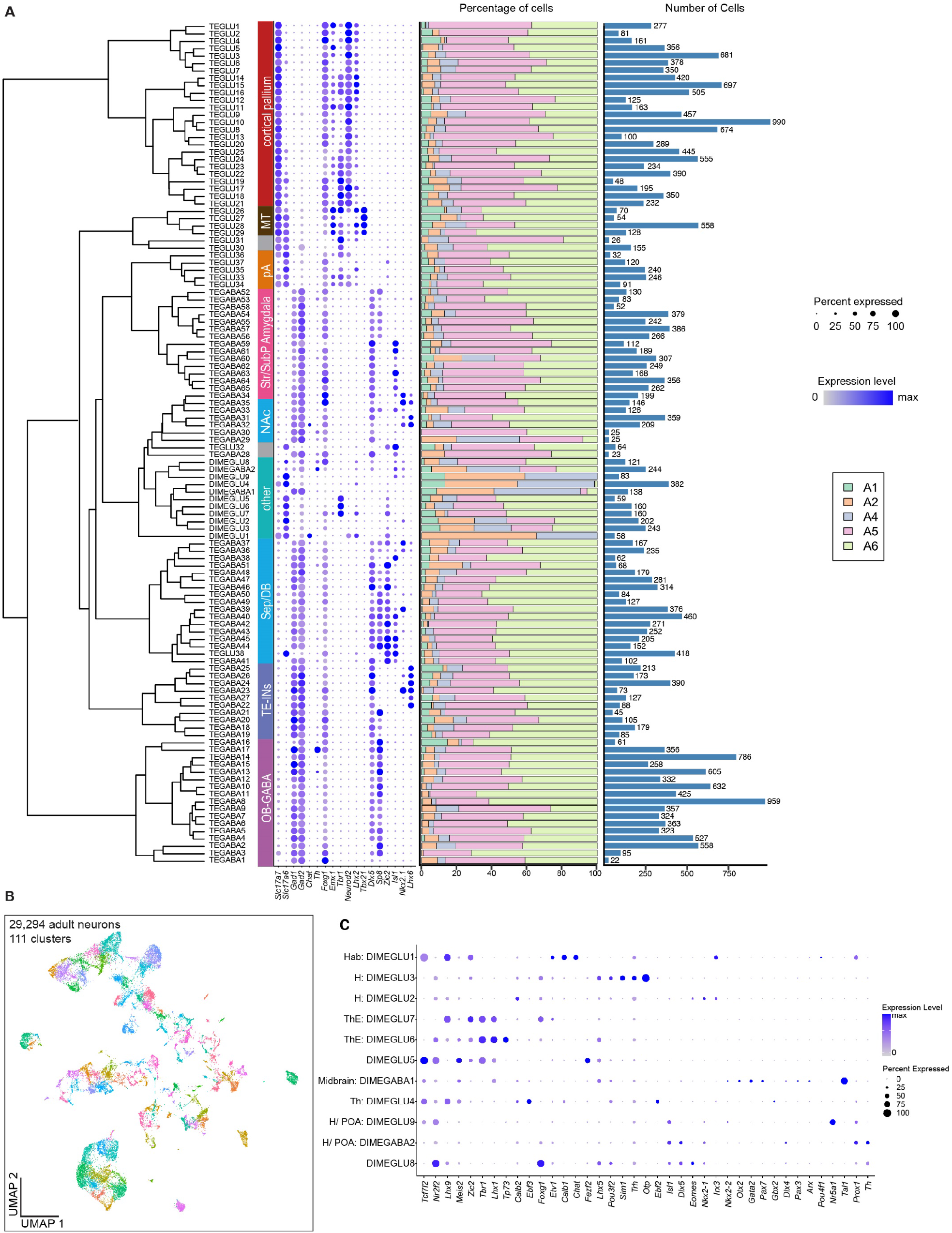
Cluster profiles and cell distribution in the adult neuronal object. **(A)** from left to right: Dendrogram of neuronal dataset, hierarchically clustered based on molecular similarity; Neuronal cluster names, corresponding to data in the same row for each plot; Dotplot of marker genes used for identification of major neuronal territories; Bar plot of percentage of cells per animal in each cluster; Bar plot of total number of cells in each cluster. **(B)** UMAP representation of 29,294 adult salamander neuron transcriptomes, colored by cluster identity **(C)** Dotplot of marker genes (columns) used for identification of neurons in the di- and mesencephalon (rows).

**Fig. S4.**
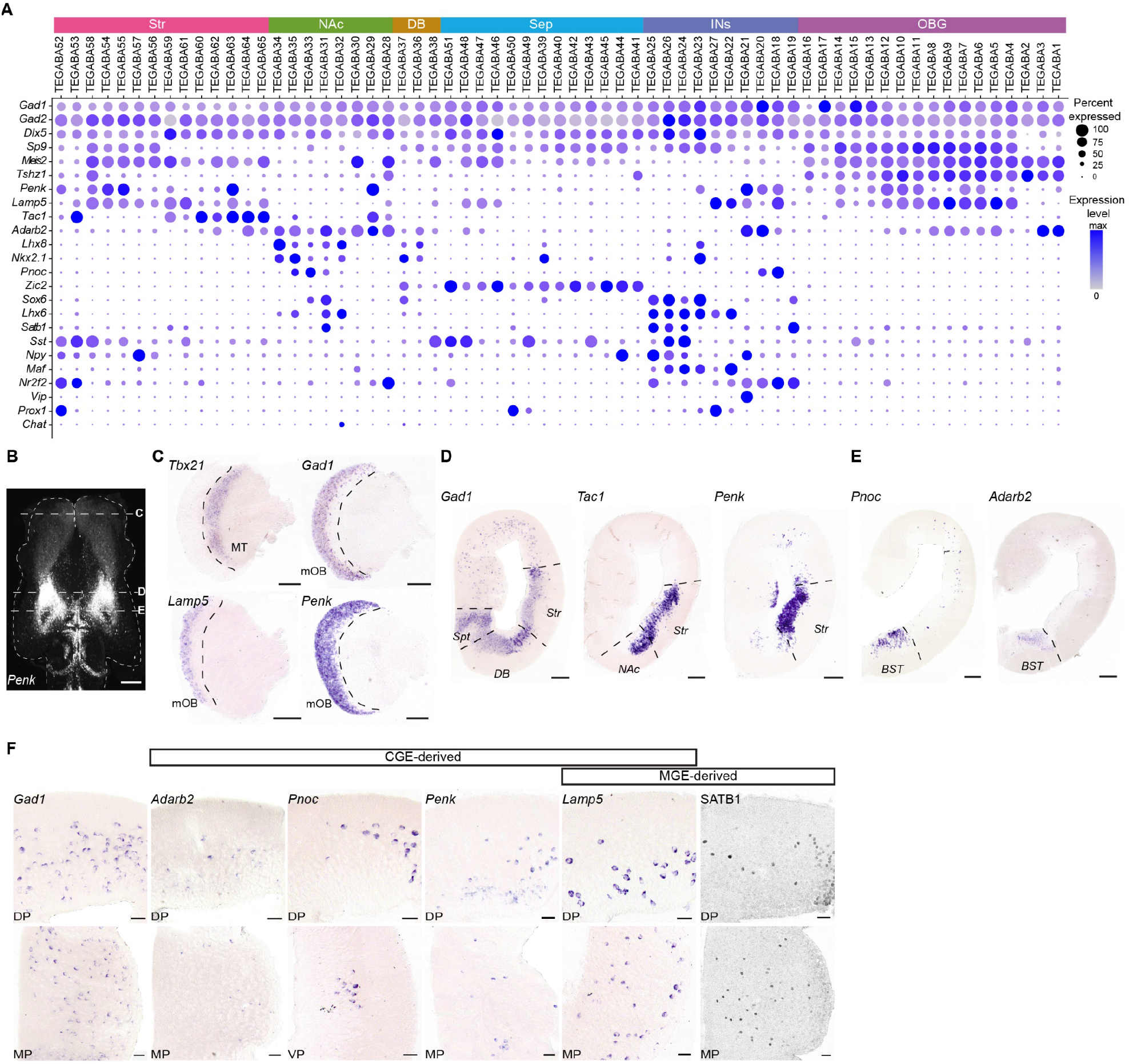
GABAergic neuron types in the *Pleurodeles* telencephalon. **(A)** Dot plot showing the expression of key marker genes defining subpallial regions and telencephalic interneurons. **(B)** Whole mount HCR-ISH for *Penk* with dashed lines indicating the sectioning plane for C-E. Scale bar: 500 um. **(C)** Coronal sections through the olfactory bulb showing mitral and tufted cells (*Tbx21+*) and olfactory bulb interneurons (*Gad1+*). Subpopulations of olfactory bulb interneurons express *Lamp5* and/or *Penk*. **(D)** Coronal sections through the telencephalon showing the expression of *Gad1* (GABAergic neurons), *Tac1* (striatum), *Penk* (striatum and subset of pallial interneurons). **(E)** Coronal sections through the telencephalon showing the expression of *Pnoc* and *Adarb2* (BST and pallial interneurons). **(F)** Magnifications from coronal sections through the telencephalon showing sparse MGE- and CGE-derived interneurons spread throughout the pallium. See methods for specifics on SATB1 antibody. Abbreviations: BST, bed nucleus of the stria terminalis; CGE, central ganglionic eminence; DB, diagonal band; DP, dorsal pallium; INs, interneurons; MGE, medial ganglionic eminence; MP, medial pallium; NAc, nucleus accumbens; OBG, olfactory bulb GABAergic; Sep, septum; Str, striatum; VP, ventral pallium.

**Fig. S5.**
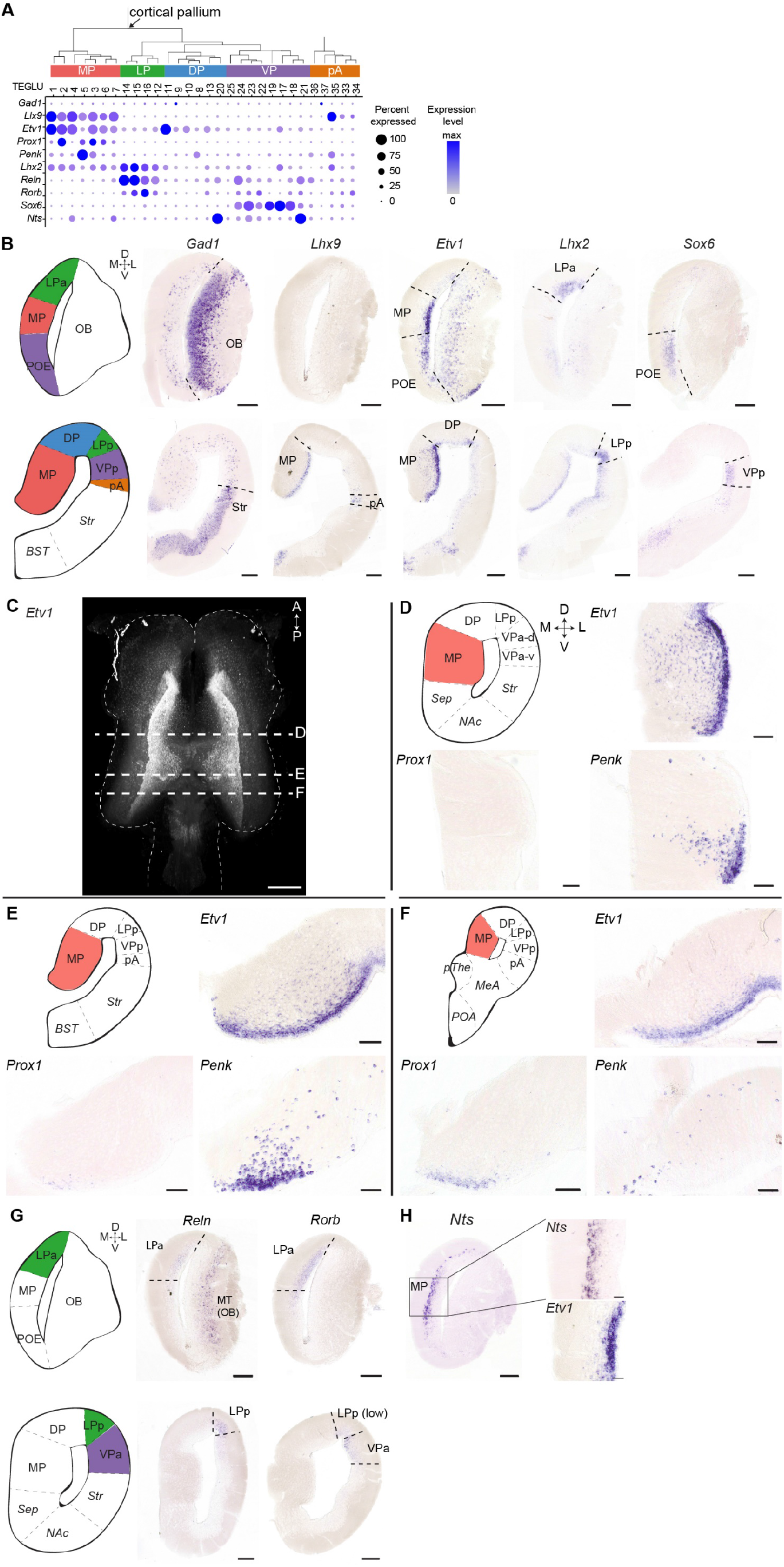
Regionalization of the pallium in *Pleurodeles*. **(A)** Hierarchical clustering and dot plot showing the expression of the genes shown in B-H. Splits in the hierarchical clustering are based on molecular similarity and do not indicate developmental or evolutionary relationships. See also Fig. S3. **(B)** Left to right: schematics of coronal sections at different anterior-posterior levels; Expression of *Gad1*, indicating the pallial-subpallial boundary, and of transcription factors labeling distinct pallial regions along the mediolateral axis: Expression of *Lhx9*, existence of a distinct rostral pallium has been proposed, posterior pallial regions include the pallial amygdala marked here; Expression of *Etv1*, the transition between medial and dorsal pallium (TEGLU 8-11, 13, 20) is clearly demarcated by a ventricular sulcus and an abrupt change of cell density; Expression of *Lhx2*, the lateral pallium is bordered by the axons of the lateral olfactory tract; Expression of Sox6, the ventral pallium is molecularly diverse, in line with its anatomical heterogeneity. Scale bars: 200 um. **(C)** Whole mount HCR-ISH for *Etv1* with dashed lines indicating the sectioning plane for D-F. Scale bar: 500 um. **(D-F)** Colorimetric ISH on floating tissue sections showing expression of *Etv1, Prox1* and *Penk* across the anterior-posterior axis in the medial pallium. *Prox1*, a marker of the dentate gyrus, overlaps with *Etv1*, a marker of the CA field, therefore showing no discrete subdivisions resembling CA or DG subfields in salamander. Scale bars: 50 um. **(G)** Left: schematics of coronal sections at different anterior-posterior levels. Right: Expression of *Reln* in the olfactory bulb mitral and tufted cells, and in the anterior and posterior lateral pallium (aLP and pLP), and of *Rorb* in the anterior and posterior LP, and in the anterior VP. Scale bars: 200 um. **(H)** Expression of *Nts* and *Etv1* in the anterior medial pallium showing the presence of at least two molecularly different layers in the pallium. Scale bars in right panels: 50um. For abbreviations, see Fig. S1; D, dorsal; L, lateral; M, medial; V, ventral.

**Fig. S6.**
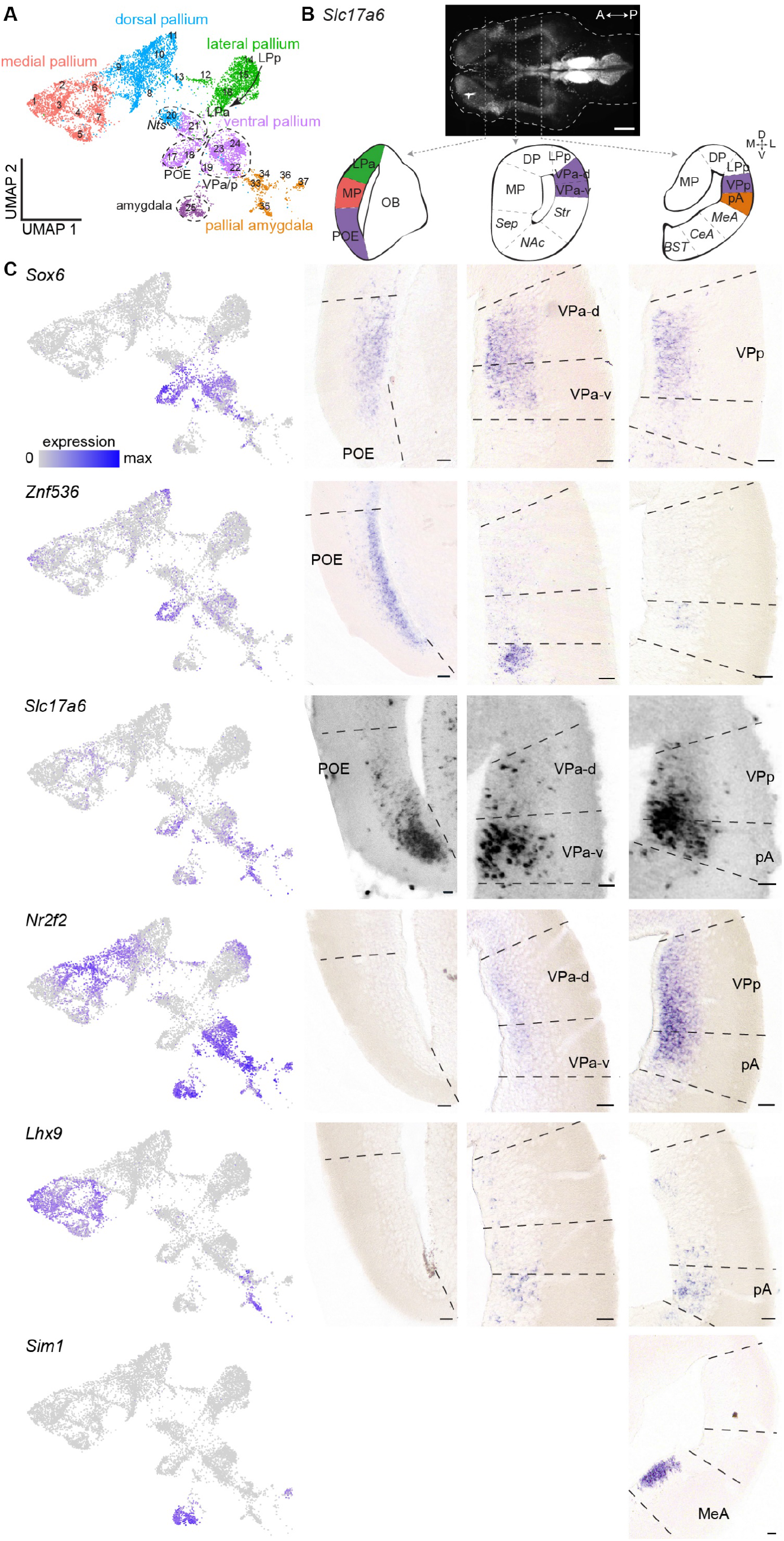
Regionalization of the ventral pallium and pallial amygdala in *Pleurodeles*. **(A)** UMAP representation of cortical pallium and pallial amygdala neurons, color coded according to region. **(B)** Whole mount HCR-ISH for *Slc17a6* and schematic overviews of the telencephalon at three levels across the anterior-posterior axis, with indication of the ventral pallium and pallial amygdala. **(C)** Left, UMAPs of pallial glutamatergic neurons, with single-cells color-coded by the expression of marker genes *Sox6, Znf536, Slc17a6, Nr2f2, Lhx9* and *Sim1*. Right, gene expression on coronal sections through the telencephalon at three representative levels across the anterior-posterior axis. These marker genes define multiple subdivisions in the ventral pallium, with an anterior *Znf536*+ POE, which is further subdivided in a more dorsal Slc17a6-negative and a ventral *Slc17a6*-positive domain. At intermediate levels, *Nr2f2* expression increases and the VP is again subdivided into a *Slc17a6*-VPa-d and a *Slc17a6*+ VPa-v. At posterior levels, ventral from the *Sox6*+/*Nr2f2*+ VPp, the pA is defined by high-level expression of *Slc17a6* and *Nr2f2*, expression of *Lhx9* in a subset of cells and absence of *Sox6*. Lastly, *Sim1*+ cells demarcate the amygdala. Scale bars: 50um. For abbreviations, see Fig. S1; D, dorsal; L, lateral; M, medial; V, ventral.

**Fig. S7.**
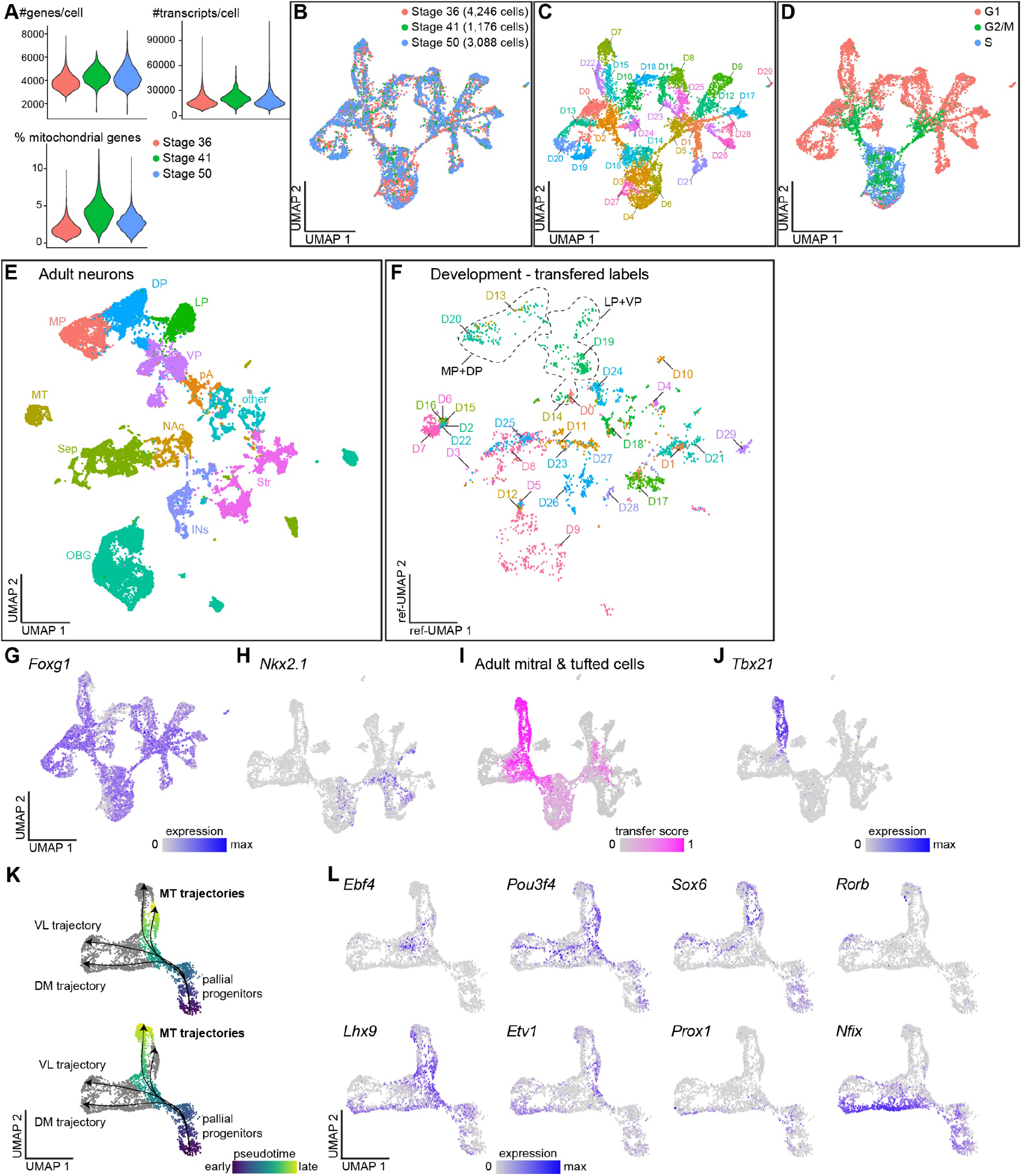
Quality control and additional data for the developmental dataset. **(A)** Violin plots of the number of genes or transcripts per cell, and percent mitochondrial genes in the full developmental neuronal dataset prior to filtering for telencephalic clusters, sorted per stage. **(B-D)** UMAP representation of 8,510 single-cell transcriptomes in the full neuronal dataset, sampled from three developmental stages, color-coded by stage (panel B), cluster (clusters D1-29, panel C) or cell cycle score (panel D). **(E)** UMAP plot of adult neurons labeled by regional identity, see also Fig. 1. **(F)** All developmental neurons that received a prediction score over 0.9 mapped on the adult UMAP space in (E), based on label transfer between adult and developmental data. Areas that correlate with adult pallial regions are demarcated by dotted lines. **(G)** Expression of *Foxg1* in the UMAP space, showing that clusters 10, 18 and 27 express no or low levels of *Foxg1* and are therefore non-telencephalic. These cluseres were removed from further analysis. **(H)** Expression of Nkx2.1 in the telencephalic developmental dataset, intended to show subpallial progenitor domain which was excluded from trajectory analysis. **(I)**Transferred cell labels from adult mitral and tufted cells onto the telencephalon developmental dataset in the UMAP space. **(J)** Expression of *Tbx21* in the UMAP space, showing that this specific marker of olfactory bulb mitral and tufted cells is expressed selectively in the olfactory bulb trajectory. **(K)** Pseudotime (Slingshot) trajectories highlighting the differentiation of mitral and tufted cells. **(L)** Expression of transcription factors differentially expressed along the dorsomedial (top) and ventrolateral (bottom) trajectories. Abbreviations: DP, dorsal pallium; DM, dorsomedial; LP, lateral pallium; INs, interneurons; MP, medial pallium; MT, mitral and tufted cells of the olfactory bulb; NAc, nucleus accumbens; OB, olfactory bulb; OBG, olfactory bulb GABAergic; pA, pallial amygdala; Sep, septum; Str, striatum; VL, ventrolateral; VP, ventral pallium

**Fig. S8.**
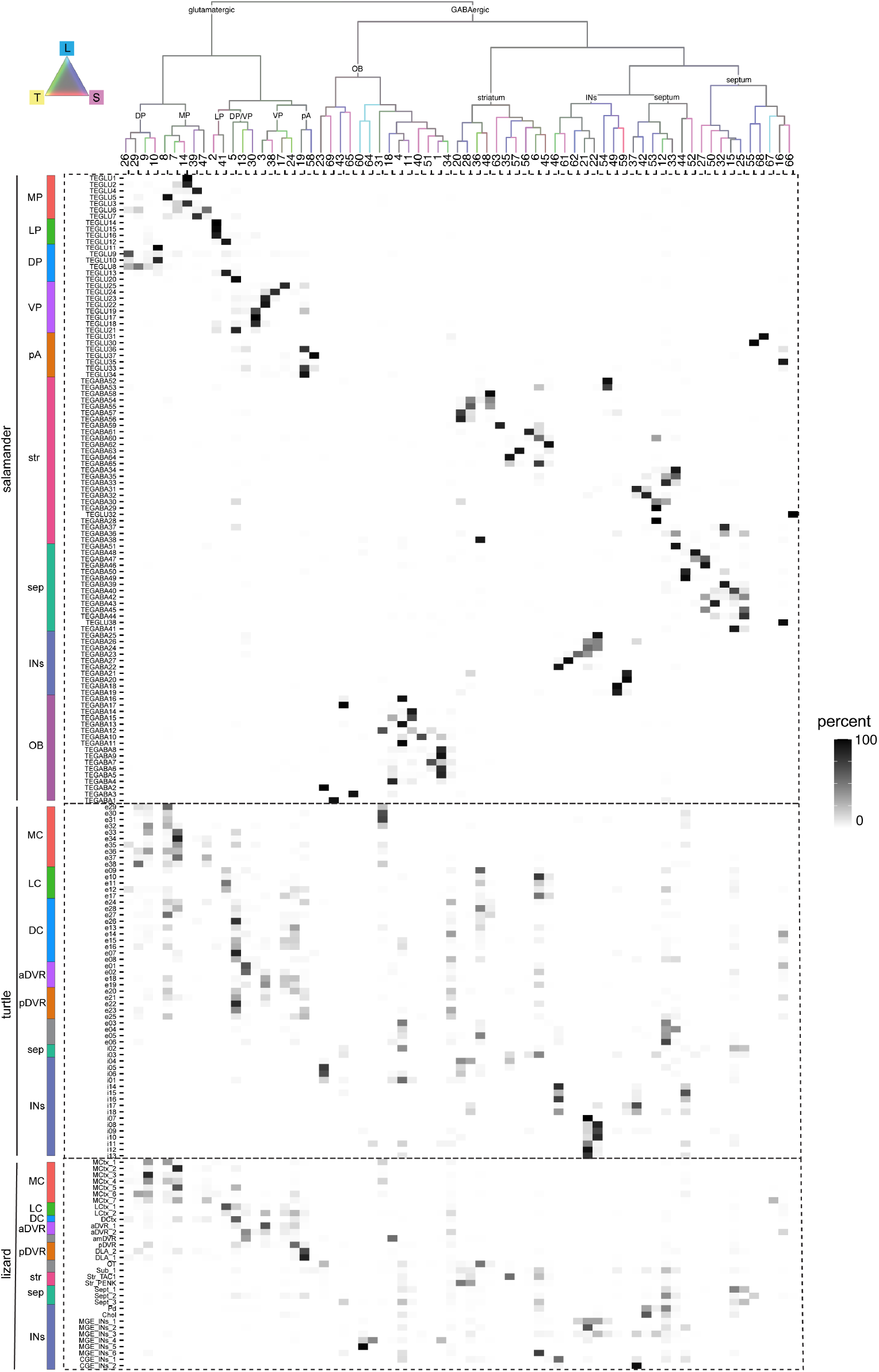
Integration of salamander, lizard, and turtle telencephalic neurons. Dendrogram (top) shows the molecular similarity of integrated clusters, with branches colored by species mixture (gray, equal proportion of cells from each species). The heatmap under the tree shows the percentage of cells from each original cell cluster according to its species-specific annotation (rows) in the integrated clusters (columns). Our integrated dataset splits broadly into two groups defined by GABAergic and glutamatergic identity.

**Fig. S9.**
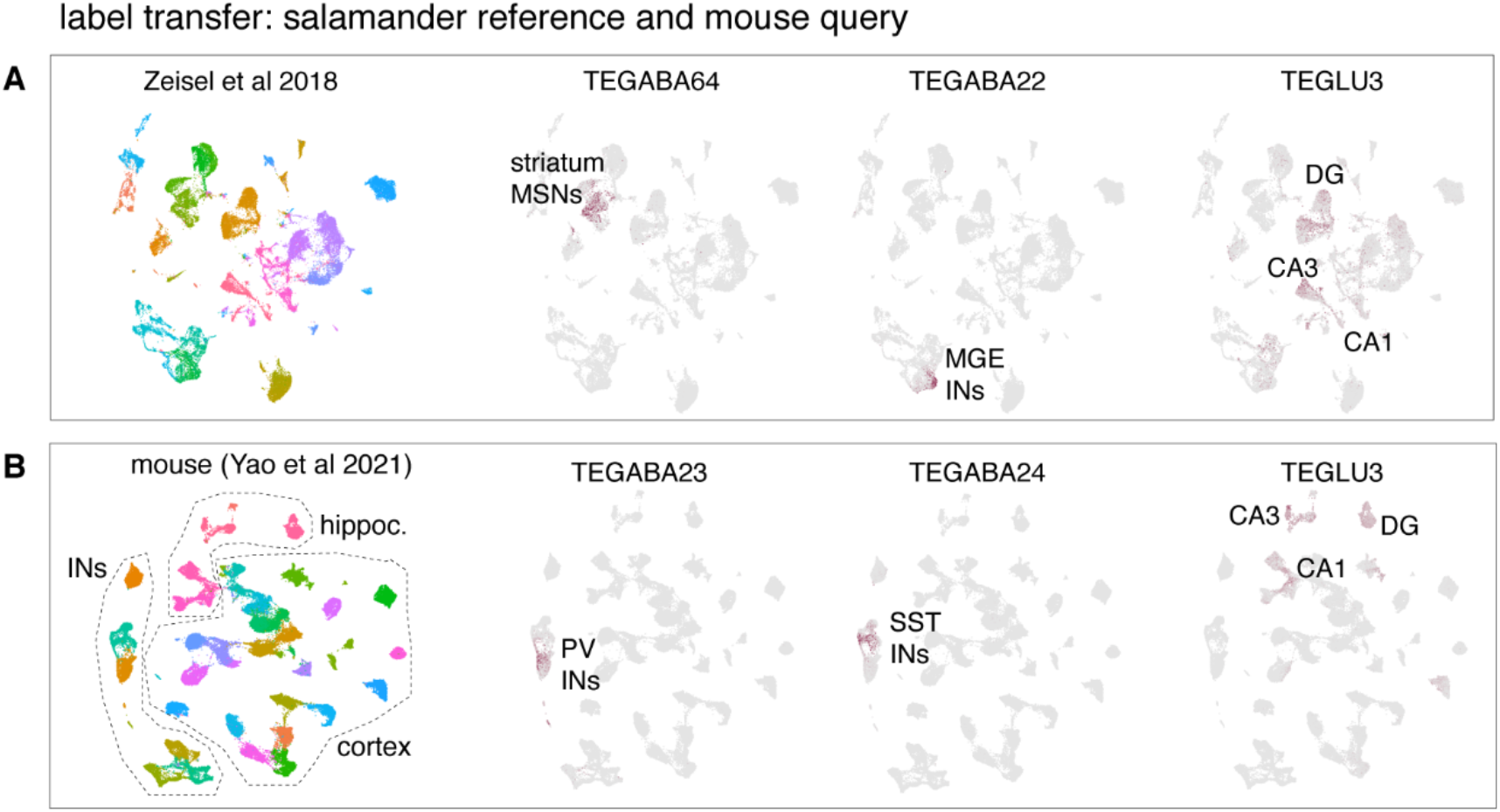
Comparison of salamander and mouse scRNAseq data by label transfer. **(A)** UMAP plots of telencephalic neurons from Zeisel et al 2018. Left: cells color-coded by cluster. Right: cells color-coded by the label transfer score for the salamander clusters indicated. Medium spiny neurons (MSN) in the mouse striatum mapped with high scores (label transfer score) on the salamander TEGABA64 cluster (striatum), mouse MGE interneurons mapped on salamander TEGABA22 cells, and mouse DG, CA1 and CA3 neurons mapped on salamander TEGLU3 cells. **(B)** Same as A, but using a downsampled Yao et al 2021 dataset (cortex and hippocampus) as query. Mouse Pvalb (PV) interneurons mapped on the salamander TEGABA23 cluster, mouse SST INs on salamander TEGABA24, and mouse DG, CA1 and CA3 neurons on salamander TEGLU3.

**Fig. S10.**
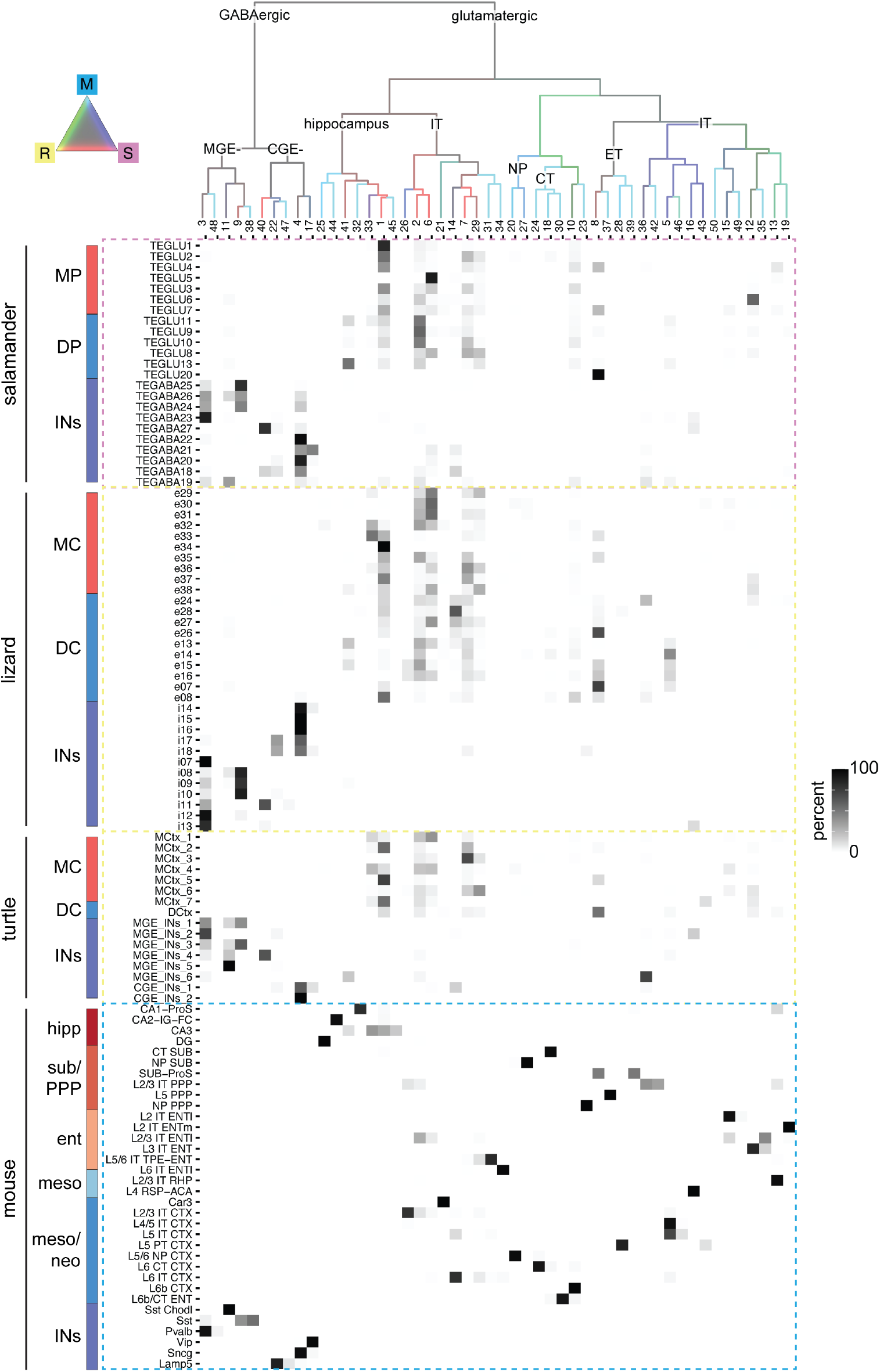
Integration of salamander, turtle, lizard, and mouse neurons from the dorsomedial pallium. Dendrogram (top) shows the molecular similarity of integrated clusters, with branches colored by species mixture (gray, equal proportion of cells from each species). The heatmap under the tree shows the percentage of cells from each original cell cluster according to its species-specific annotation (rows) in the integrated clusters (columns).

**Fig. S11.**
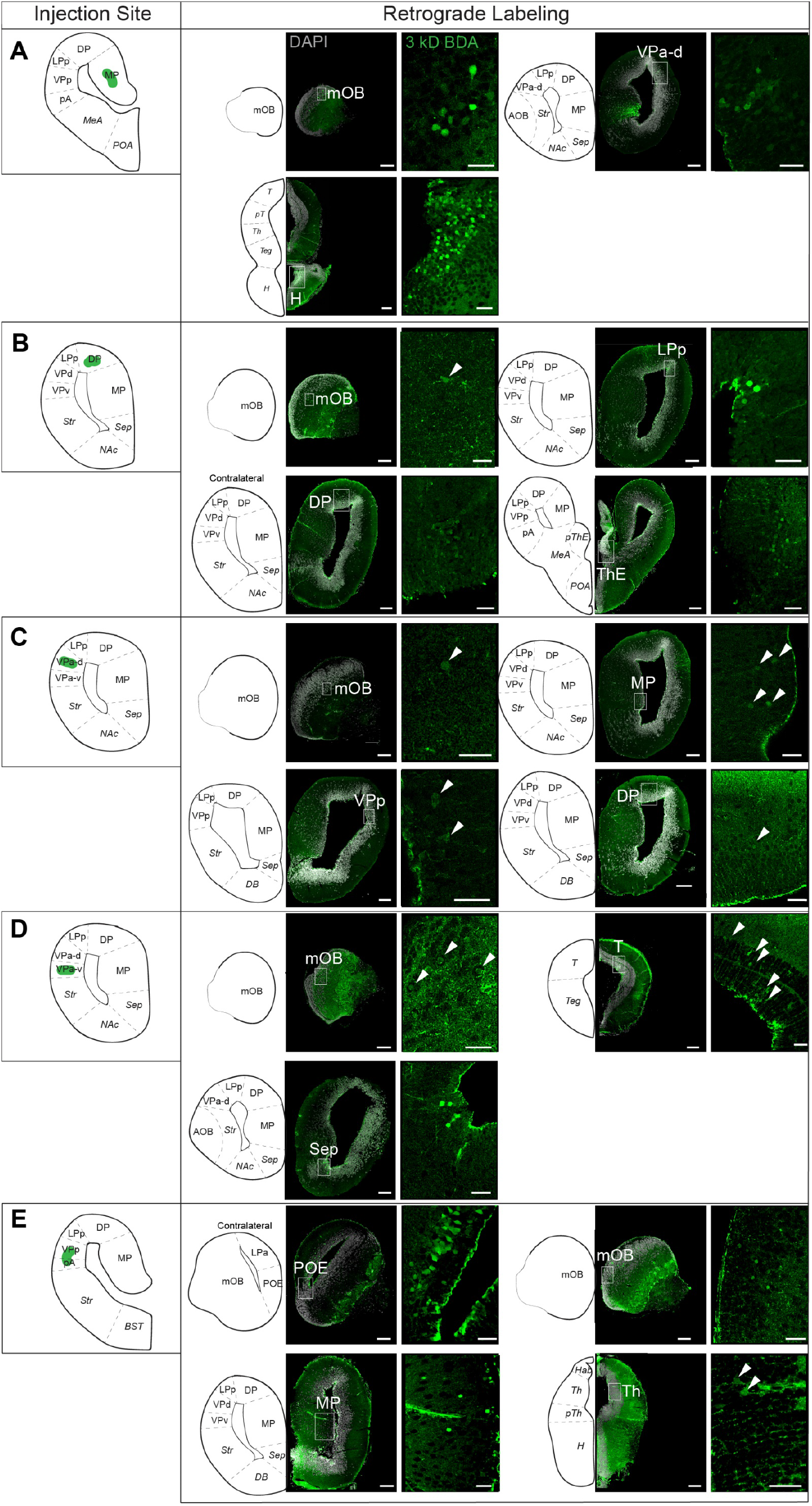
Additional tracing data: Pallial injections. Additional representative images of 3 kD BDA tracer injections and retrograde labeling into **(A)** MP **(B)** DP **(C)** VPa-d **(D)** VPa-v **(E)** VPp/pA. For neuroanatomical abbreviations, see Fig. S1. Scale bars: 200 um for slice overview; 50 um for zoomed images

**Fig S12.**
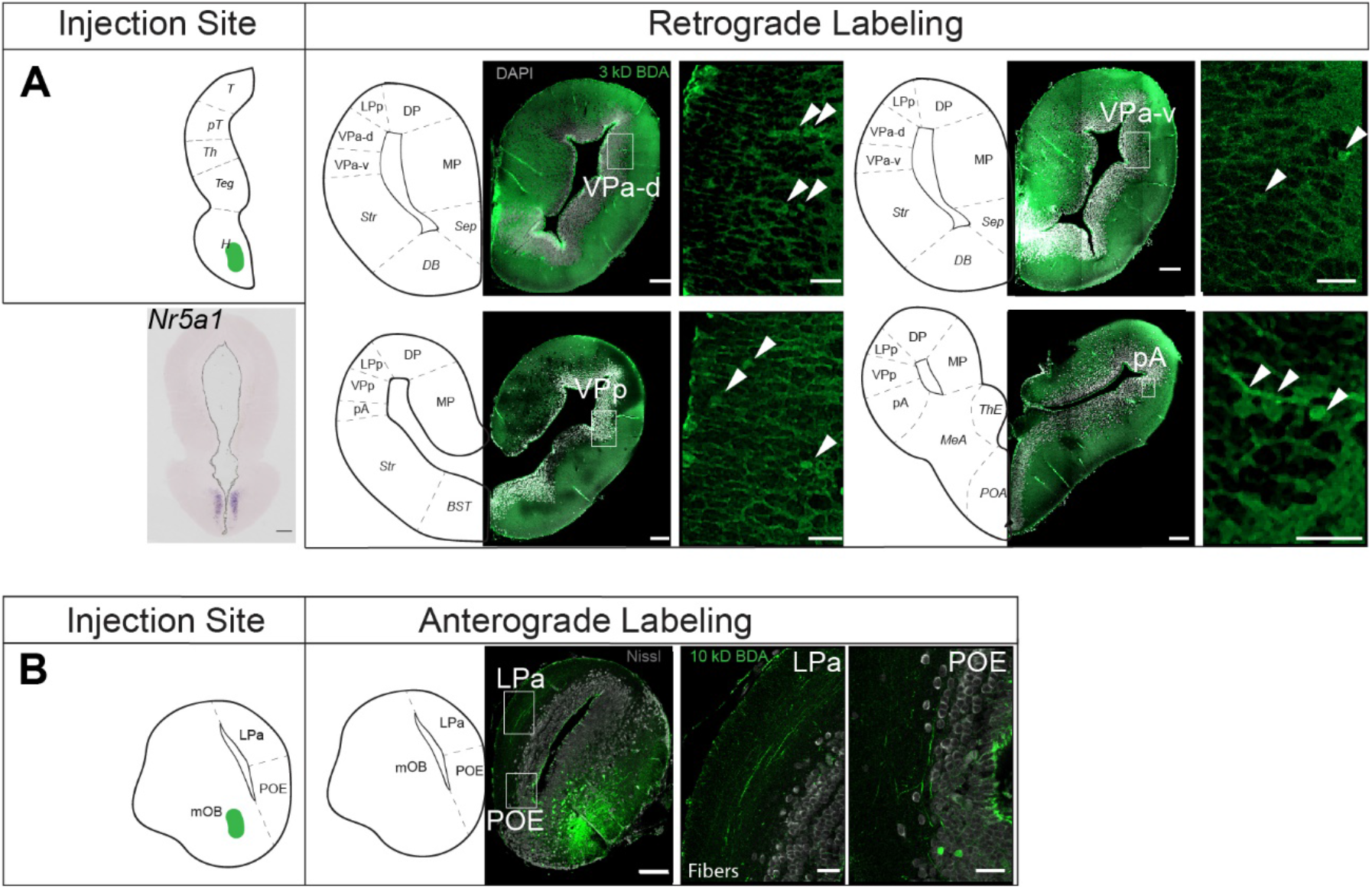
Additional tracing data: Extra-pallial injections. Additional representative images of BDA tracer injections and labeling of **(A)** VMH injection and retrogradely labeled cells; **(B)** mitral tufted cells of the mOB and anterogradely labeled fibers projecting to pallium. Scale bars: 200 um for slice overview; 50 um for zoomed images

